# Intestinal stem cells enhance local mucosal immunity through apoptotic body phagocytosis

**DOI:** 10.1101/2024.10.29.620856

**Authors:** Julie Y. Zhou, Qiuhe Lu, Yujia Hu, Satoru Fujii, Scott T. Espenschied, Morgan J. Engelhart, Kelsey J. Lewis, Paul E. Karell, Yi Han, Heaji Shin, Robert E. Schmidt, Daniel J. Silver, Andrei I. Ivanov, Omer H. Yilmaz, Thaddeus S. Stappenbeck

## Abstract

Modulation of immune tone at mucosal surfaces is critical to maintain homeostasis while facilitating the handling of emerging threats. One dynamic component of immune modulation is the phagocytosis and clearance of apoptotic bodies known as efferocytosis that inhibits inflammation by promoting its resolution. Here, we evaluated the effects of apoptotic body phagocytosis by intestinal epithelial stem and progenitor cells (ISCs). Unexpectedly, instead of immunomodulation through efferocytosis, this process elevated local immune system activity. To achieve this result, ISCs actively engaged apoptotic bodies in a unique fashion, leading to their engulfment and ultimate delivery to lysosomes for processing. We found that ISCs were capable of actively recruiting inert material such as apoptotic bodies by using actin-based intrinsic biomechanical processes. Uptake of apoptotic bodies was facilitated by complement factor C3 produced by apoptotic bodies themselves. ISCs in turn generated signals heightening T cell activity that was driven in part by ISC-generated TNF. Taken together, uptake of apoptotic bodies by ISCs produced a local inflammatory alert to specific immune cells. This altered paradigm for the response to phagocytosed apoptotic bodies fits the needs of active mucosal surfaces and demonstrates that efferocytosis as currently defined is not a universal response of all cell types.

## INTRODUCTION

The innermost layer of the mammalian intestine, the mucosa, is lined by a single layer of epithelial cells which constitute a selectively permeable barrier. The rapid and continuous renewal of this barrier is driven by multipotent intestinal epithelial stem and progenitor cells (ISCs) located in the crypts of Lieberkühn^1,2^. During phases of damage and subsequent repair, ISC proliferative activity and production of specific cell lineages can be dynamically altered. Given the critical role of the epithelial barrier in limiting translocation of luminal microbes and their products, it has long been hypothesized that ISCs could act as sentries, transducing signals to the mucosal immune system to clear infections or repair damage. However, several studies of primary ISCs indicate that they have low to non-detectable expression of key innate receptors that would mediate this function^3,4^. The expression of such innate receptors is well known to be abundant and functional in innate immune cells within the intestine. Signals from these cells in turn provide cues to ISCs to adjust lineage allocation and turnover rate in response to environmental perturbations^5–8^. The limitation of this primarily unidirectional communication model is that it relies on barrier disruption to activate immune responses, which may occur too late to contain damage effectively. This raises the question of whether ISCs can directly sense the health of their niche and alert the local immune system earlier. Recent findings revived the idea that ISCs may engage with mucosal T cells by acting as non-professional antigen-presenting cells through the inducible expression of MHC class II^7,9–11^. While these studies highlight ISC involvement in immune crosstalk, whether ISCs can actively initiate such interactions remains unknown.

In searching for a modality by which ISCs can monitor their environment, we revisited work from the 1970s, when Hazel Chang and Charles Leblond first provided evidence for the multipotency of ISCs through observing engulfment of apoptotic bodies by crypt base columnar cells. They traced the apoptotic body remnants into differentiated progeny that included all major lineages of the intestinal epithelium^12^. While the phagocytic ability of ISCs functionally served as an early lineage tracing tool, many questions remained: how do ISCs uptake apoptotic cells, what biological sequelae ensue, and why do ISCs perform this task?^13^ Recently, several groups have demonstrated that stem cells residing in niches that undergo cyclic developmental waves of cell death can engage in a process characterized by non-inflammatory efferocytosis, including hair follicle stem cells during the catagen cycle of this structure^14,15^, migrating neural crest stem cells^16^, and cells of the developing extraembryonic gut^17^. Other studies have also shown that differentiated intestinal epithelial cells (IECs) including enterocytes and Paneth cells can clear apoptotic cells^18–20^. These studies all support a model in which epithelial cells, including stem cells, can perform the clearance function of macrophages under specific circumstances, although the biological significance of this activity beyond local debris clearance is not well understood.

Here, we propose that apoptotic body engulfment by ISCs serves as a sensor for early damage in the intestine and is involved in local intercellular communication, challenging the traditional view that tissue-resident adaptive immune cells are activated via myeloid cells through specific modalities only after epithelial barrier compromise. We recapitulated the work of Cheng and LeBlond using modern reagents and found that this process of damage leads to immune enhancement, especially of T cells in the mucosa. Using *in vitro* methods to cultivate primary ISCs under specific conditions^21–23^, we explored mechanisms of recruitment, engulfment, and intracellular processing of apoptotic bodies and factors that can activate the local immune system. Leveraging strategies inspired by work performed by pioneers in the 1970s, we modeled cell death in the intestinal crypts in mice and found local immune stimulation. We developed a model system to study the mechanisms underlying ISC engulfment of apoptotic epithelial cells and the immunologic consequences of this activity.

## RESULTS

### Enhanced local intestinal immune activity occurs with crypt-epithelial cell engulfment of apoptotic bodies

Differentiated epithelial cells that line villi and surface cuffs in the small intestine and colon, respectively, typically undergo apoptosis and are removed through luminal extrusion. Conversely, epithelial cells that undergo apoptosis within crypts can be retained within the crypt epithelium where previous work suggests their engulfment by neighboring cells including stem cells^12^. To examine the consequences of this activity in a mouse experimental system, we leveraged the known properties of the thymidine analog, 5-ethynyl-2-deoxyuridine (EdU), to arrest cell cycle and induce cell death (predominately apoptosis) in a subset of intestinal epithelial stem and progenitor cells (ISCs) (Fig. 1a)^24^. Pulsing with a single dose of EdU that effectively targets dividing cells,^25^ we observed an increase in apoptotic bodies distributed throughout intestinal crypts within 24 hours of administration (Extended Data Fig. 1a-b). This result suggested that they did not undergo efficient luminal extrusion or clearance by professional phagocytes. To model a persistent increase in crypt epithelial apoptosis, we administered EdU (T_1/2_ = ~30 min^26^) every other day (q.o.d.) for seven days to allow for multiple waves of apoptotic cell generation and to enhance downstream signals from this process. This regiment resulted in sustained elevated numbers of EdU^+^, cleaved caspase 3 (cC3)^+^ apoptotic bodies in the crypt epithelium (Extended Data Fig. 1c). This signal was primarily localized to intestinal epithelial crypts with very limited labeling of cells in secondary immune tissues such as Peyer’s patches (Extended Data Fig. 1d). EdU signal was often contained by large (>3μm) LAMP1^+^ lysosomes within unlabeled epithelial cells of the crypt (Fig. 1b-c, Extended Data Fig. 1e) that appeared following EdU exposure (Fig. 1d). These results suggest crypt epithelial cells can contribute to apoptotic body clearance by engulfment and delivery to lysosomes.

**Fig. 1:**
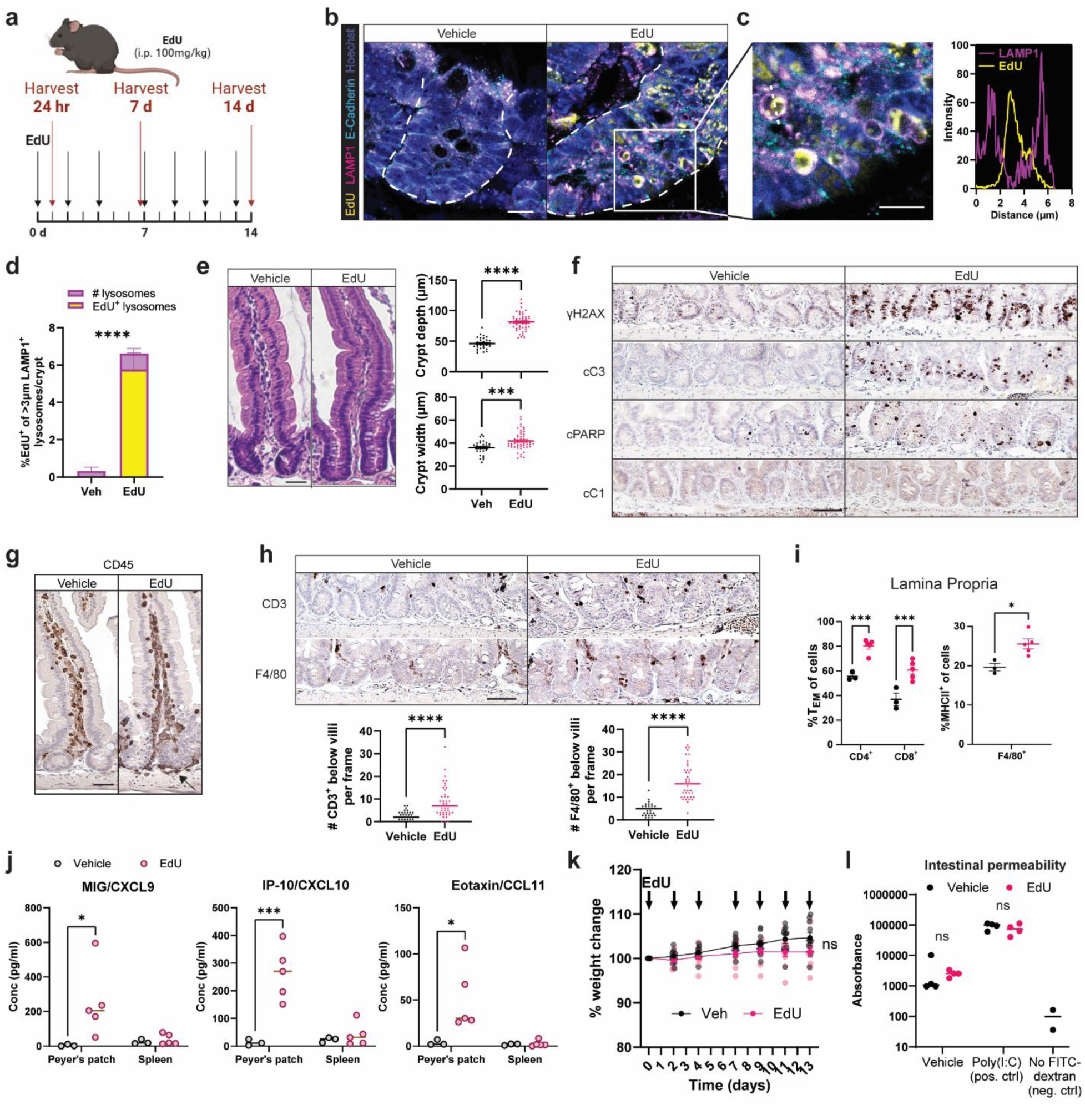
Enhanced local intestinal immune activity occurs with crypt-epithelial cell engulfment of apoptotic bodies. **a**, Model of i.p. EdU mouse experiments. **b**,**c**,**d**,**e**, Mice were injected q.o.d with EdU for 7 d. *n* = 3-5 mice per treatment group. Representative sections of proximal small intestine stained for indicated targets with (**b**) white dotted lines denoting crypts. Scale bar = 50 µm. (**c**) Magnification of left inset with white dotted line indicating intensity of LAMP1 and EdU calculations. Scale bar = 10 µm. (**d**) Quantification of large (>3 µm) lysosomes with % EdU^+^ in yellow. (**e**), Representative sections of proximal small intestine stained by H&E. Scale bar = 10 µm with quantification of crypt depth and crypt width, *n =* 30-50 crypts per treatment group evaluated. **f**,**g**,**h**, Mice were injected q.o.d with EdU for 7 days. *n* = 3-5 mice per treatment group. Representative sections of proximal small intestine. Scale bars = 50 µm. Staining for (**f**) γH2AX, cleaved caspase 3 (cC3), cleaved PARP (cPARP), and cleaved caspase 1 (cC1). (**g**) CD45, (**h**) CD3 and F4/80 with (Bottom) showing quantification of stained cells per frame. n = 30- 50 frames per condition. **i**, (Left) Frequency of T effector memory (T_EM_, CD62L^-^ CD44^+^) within the indicated T cell compartment and (Right) frequency of MHCII^+^ events of F4/80^+^ cells of lamina propria in mice treated or untreated with 14 d EdU. *n* = 3-5 mice per genotype. **j**, Cytokine analysis of cells isolated from the Peyer’s patch or spleens of mice treated with EdU for 14 d. *n* = 3-5 mice per treatment group. **k**, Weight change of mice over the course of 14d with arrows indicating EdU administration. *n* = 10 mice per condition. **l**, Intestinal permeability following 14d of EdU administration as assessed by FITC-dextran gavage and detection of FITC signal in serum. Poly(I:C) was used as a positive control for intestinal damage. *n* = 4 for each condition (n = 2 for no FITC control). Data are representative of at least 2 independent experiments. One-way analysis of variance (ANOVA) with Tukey’s multiple comparisons post hoc test was performed (**i**,**j**,**k**,**l**); Two-tailed Student’s t test in (**d**,**e**,**h**). *p < 0.05, **p < 0.01, ***p < 0.001, ****p < 0.0001. ns, not significant.

Interestingly, we found that a 1-week course of EdU exposure led to crypt hyperproliferation throughout the small intestine, demonstrated by increased crypt depth (Fig. 1e, Extended Data Fig. 1f), as well as lineage allocation skewing towards absorptive cells in the colon (Extended Data Fig. 1f). These injury-associated ISC behaviors suggested that apoptotic cell engulfment by ISCs led to activation of components of the immune system^27,28^. In support of this, we found that after treating mice with EdU for 2 weeks, enrichment of cC3, cPARP, γH2AX immunoreactivity [with no detectable increase in the pyroptotic marker cleaved caspase 1 (cC1)] suggested cell death in the crypt was predominantly apoptotic (Fig. 1f). Notably, there was increased CD45^+^ staining near crypts (Fig. 1g) and macrophages with an activated morphology interspersed with CD3^+^ T cells enriched around crypts (Fig. 1h) in EdU-treated mice. Flow cytometric analysis revealed that T cells were skewed towards a T_EM_ phenotype (CD44^+^CD62L^-^) and F4/80^+^ macrophages adopted an inflammatory phenotype (upregulated expression of MHCII) in the lamina propria (Fig. 1i, Extended Data Fig. 2a) but not in the Peyer’s patch (Extended Data Fig. 2b), mesenteric lymph node (mLN) (Extended Data Fig. 2c), or spleen (Extended Data Fig. 2d). Cytokine analysis revealed an increase in chemokines secreted by local lymphoid cells but not splenic cells (Fig. 1j). These results indicate that the enhanced immunity in response to crypt apoptosis was local and not a systemic phenomenon. While excess cell death in intestinal crypts led to measurable immune enhancement in select cell types, it did not elicit uncontrolled inflammation either within the local tissue or in distant sites. Importantly, mice maintained overall weight (Fig. 1k) and epithelial barrier function (Fig. 1l) during the 2-week exposure to EdU.

Taken together, these data suggest that epithelial cells in the intestinal crypts engulf apoptotic bodies. This leads to targeted immune enhancement in local tissues, particularly in the proximal small intestine with limited PAMPs, suggesting a cell intrinsic mechanism. This model of immune enhancement originating from the epithelium to the mucosal immune system is not in concordance with the current model that uptake of apoptotic bodies by any stem cell would lead to efferocytosis and downmodulate immune responses^14,29^. Our model is that sustained apoptotic body uptake by ISCs is the danger signal that affords this type of stem cell the opportunity to locally fortify immune defenses without creating an uncontrolled inflammatory response that could damage the tissue. Therefore, we sought to understand the biology underlying this process and why ISCs have a countervailing response to apoptotic body uptake.

### ISCs along the intestinal tract uptake apoptotic iIECs

We used a reductionist approach to test whether ISCs in isolation can engage in this behavior in response to apoptotic body uptake. We used intestinal epithelial spheroid culture derived from intestinal crypts that were enriched for stem and progenitor cells^30^. We modeled epithelial cell uptake of exogenous apoptotic bodies by supplementing fluorescently labeled apoptotic immortalized intestinal epithelial cells (iIECs treated with staurosporine) to 3D mouse intestinal spheroid cultures that are enriched in ISCs (Fig. 2a). To increase the number of cells available for treatment, we used an immortalized cell line derived from primary mouse intestinal epithelial cells (Extended Data Figs. 3a-f). Three days after co-incubation, we observed apoptotic iIECs within ISCs (Fig. 2b), consistent with engulfment; these engulfed apoptotic iIECs were surrounded by the actin cytoskeleton of ISCs (Fig. 2c, see supplemental video 1 for 3D reconstruction). This property was generalizable to ISCs derived from all segments of the mouse intestine; apoptotic bodies labeled ~18% of ISCs after 3 days of co-incubation when added in a 1:1 ratio (Fig. 2d, Extended Data Fig. 4a-c). Titrating the number of apoptotic iIECs added to ISCs demonstrated that uptake occurred in a dose-dependent manner with a maximum of 30% labeled ISCs even when cultures were densely populated with apoptotic iIECs (5:1 ratio of apoptotic iIECs to ISCs) (Fig. 2e). Time course experiments with a 1:1 ratio (apoptotic iIEC to ISCs) showed that labeled apoptotic iIECs progressively accumulated in ISCs during the course of the experiment (1-3 days), suggesting that uptake outpaced degradation (Fig. 2f, Extended Data Fig. 4d). To confirm that spheroids were not limited to engulfing apoptotic iIECs, we supplemented apoptotic primary and immortalized cells from multiple sources to ISCs, all of which were similarly engulfed (Extended Data Fig. 4e).

**Fig. 2:**
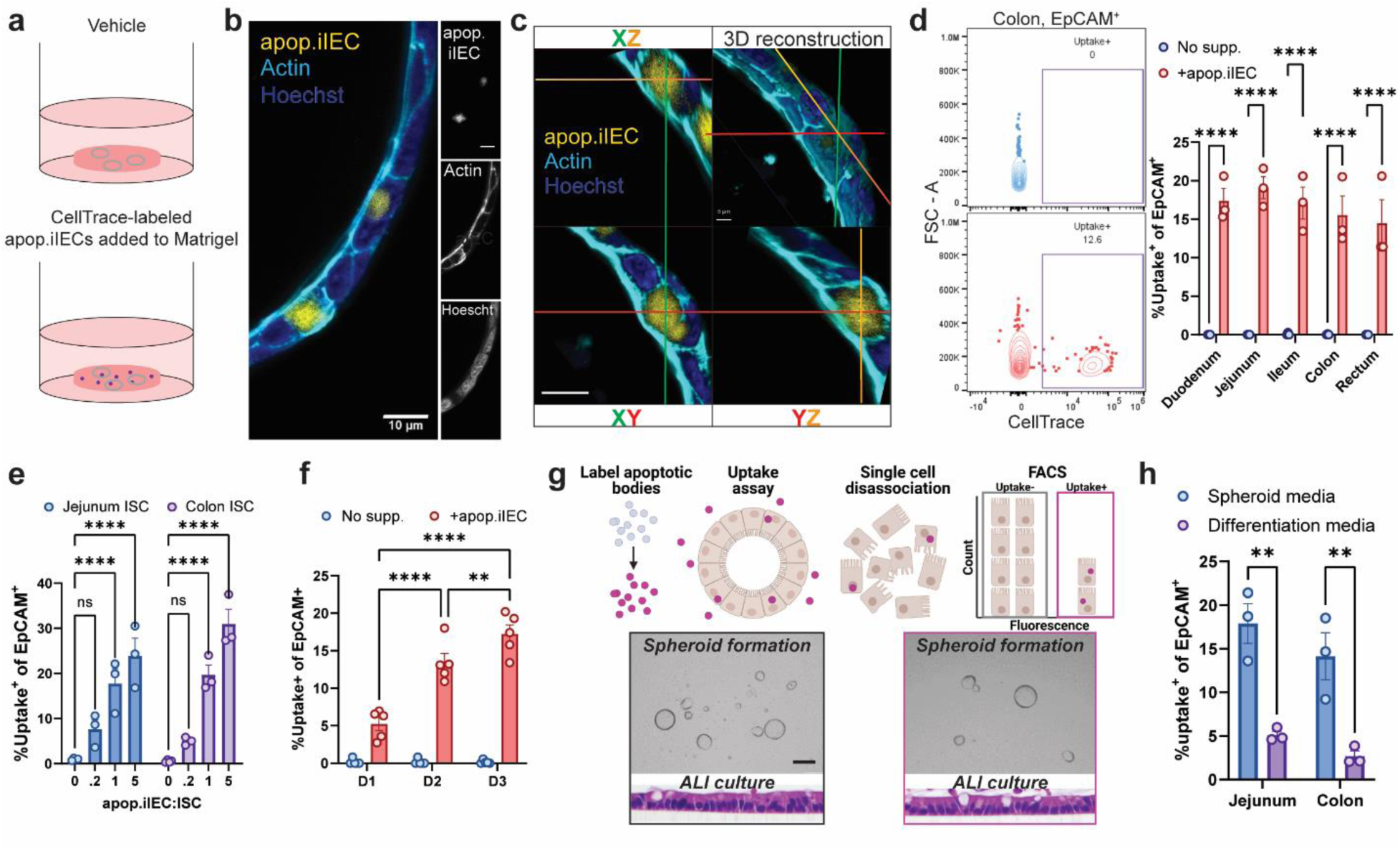
ISCs along the intestinal tract uptake apoptotic iIECs. **a**, Schematic of ISC spheroid culture supplemented with fluorescently labeled apoptotic immortalized intestinal epithelial cells (apop.iIECs). **b**, (left) Merged channels of an immunofluorescent-stained ISC spheroid supplemented with CellTrace-labeled apop.iIECs. (right) individual channels. Scale bar = 10 µm. *n* = 3 replicates. **c**, 3-dimensional reconstruction of an ISC spheroid with cross sections in XZ, XY, and YZ planes. Scale bar = 10 µm. *n* = 3 replicates. **d**, Flow cytometric analysis of the frequency of uptake^+^ ISCs (Sytox^-^EpCAM^+^CellTrace^+^); (left) representative histogram and (right) quantification of % uptake^+^ of EpCAM^+^ ISCs grown from crypts harvested along the intestinal tract and incubated with CellTrace-labeled apop.iIECs. *n* = 3 replicates. **e**, Flow cytometric analysis of the frequency of uptake^+^ ISCs (from jejunum or colon) with the indicated ratios of apop.iIEC to ISCs. The number of ISCs were held constant. *n* = 3 replicates. **f**, Flow cytometric analysis of the frequency of uptake^+^ ISCs after the indicated time points. *n* = 5 per time point. **g**, Spheroid formation and air-liquid-interface epithelial differentiation assays (H&E) of FACS purification of uptake^+^ or uptake^-^ ISCs. *n* = 3. **h**, Flow cytometric analysis of the frequency of uptake^+^ ISCs following culture in spheroid media or differentiation media. *n* = 3. Data are representative of at least 2 independent experiments. One-way analysis of variance (ANOVA) with Tukey’s multiple comparisons post hoc test (**d**, **e**, **f**, **h**); **p < 0.01, ***p < 0.001, ****p < 0.0001. ns, not significant.

To confirm the stem and progenitor function of engulfer cells, we tested FACS-purified uptake^+^ and uptake^-^ cell behavior *in vitro*. We found that uptake^+^ cells generated spheroids when seeded in 3D culture, suggesting self-renewal capability. In addition, spheroids grown from uptake^+^ ISCs differentiated into multi-lineage epithelium when exposed to air-liquid interface (ALI) conditions^31^ (Fig. 2g). Therefore, uptake^+^ cells satisfied the functional definition of stem/progenitor cells by 1) exhibiting the ability to self-renew and 2) give rise to multiple differentiated daughter cell lineages. Further, by blocking EP4 to induce differentiation^22^, we observed that differentiated cells had a reduced ability to uptake apoptotic iIECs (Fig. 2h). Altogether, these data supported historical findings that ISCs from along the intestinal tract can uptake apoptotic bodies and established a working model of this system using 3D spheroid culturing techniques.

### Mechanical forces generated by actin-mediated membrane rearrangement drive uptake of apoptotic iIECs

Given that ISCs were able to engulf apoptotic bodies, we next sought to determine how spheroids engage immobile apoptotic bodies. We hypothesized two non-exclusive models, directional migration and recruitment, that could precede engulfment (Fig. 3a). Using time-lapse live cell imaging, directional migration was observed in spheroids cultured with labeled apoptotic iIECs added immediately after passage; here we used live cell imaging to show that spheroids migrated towards apoptotic iIECs over distances that could exceed 100 µm over 5 hours (or approximately 20 µm/hr)(Fig. 3b, Supplemental video 2). Direct recruitment was most prominent when adding apoptotic iIECs to established (2-day old) spheroid cultures: here, spheroids were capable of drawing in many apoptotic iIECs or beads from multiple directions, simultaneously (Fig. 3c and Supplemental video 3 for beads, Supplemental video 4 for apoptotic iIECs). Recordings including both of these behaviors were subsequently used to train the machine learning-based user-defined behavior tool, LabGym^32,33^ (Fig 3d), which enabled reliable classification and quantification of recruitment, migration, and stationary (no directed movement, recruitment, or migration) behaviors (Fig. 3e). Remarkably, 80% of spheroids exhibited recruitment activity when grown in ISC media, however recruitment ability was lost when spheroids were cultivated in differentiation media (Fig. 3e, Supplemental video 5), consistent with reduced uptake by differentiated enterocytes (Fig. 1h).

**Fig. 3:**
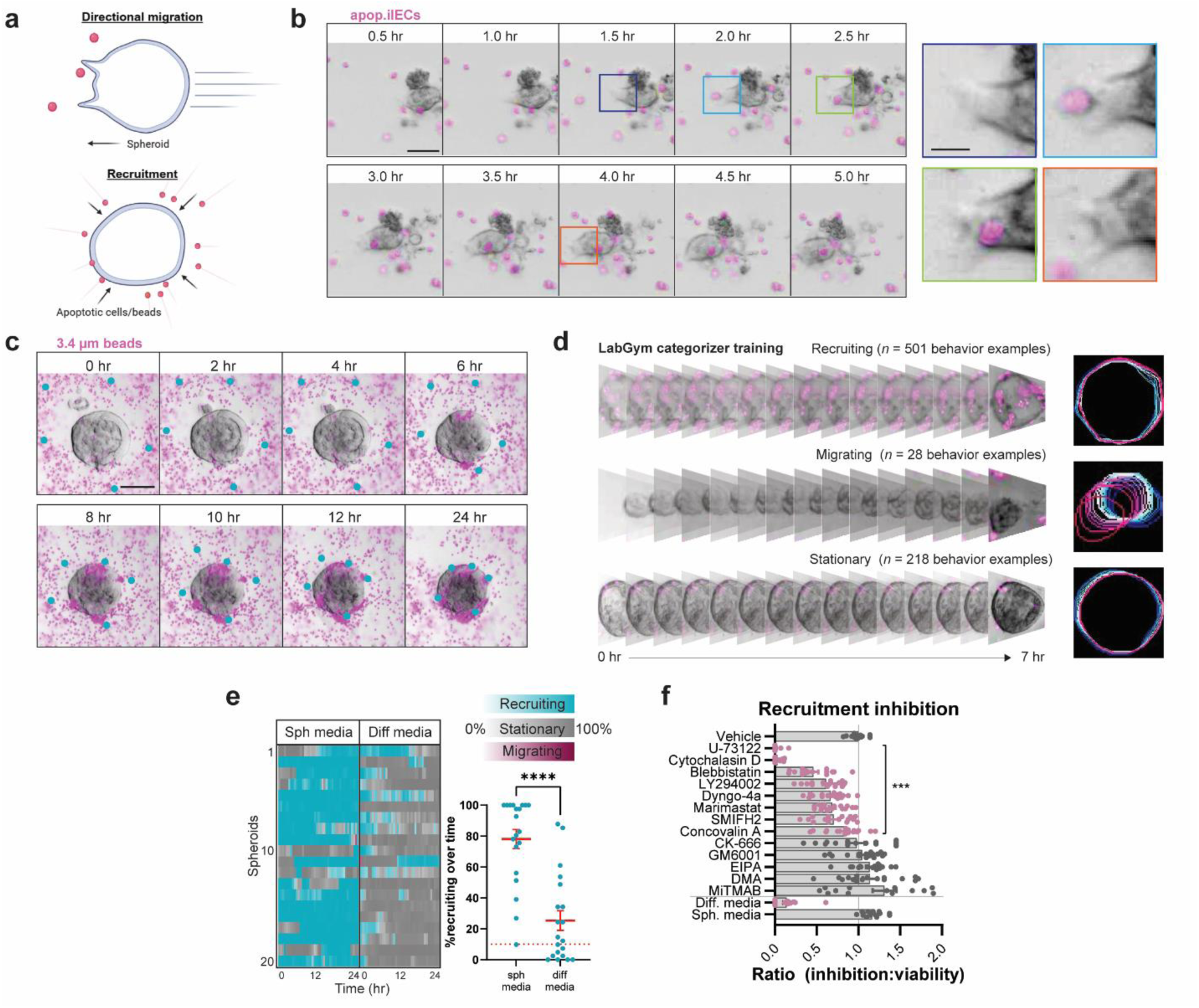
Mechanical forces generated by actin-mediated membrane rearrangement drive uptake of apoptotic iIECs. **a**, Schematic of the proposed mechanisms underlying ISC and apoptotic body proximity. **b**, Spheroid and apop.iIEC (labeled magenta) culture imaged at the indicated time points. Left: scale bar = 50 µm. Right: box color denotes origin of inset. Scale bar = 20 µm. **c**, Spheroid and bead (labeled magenta) culture imaged at the indicated time points. Turquoise color was added *post hoc* to indicate the location of specific beads. Scale bar = 50 µm. **d**, Example of spheroid behavior training in LabGym following generation of subject traces. Each frame and trace represent 30 min of live imaging. **e**, Behavior categorization timeline of Recruiting, Stationary, and Migrating behaviors of spheroids cultured in spheroid (sph) medium or differentiation (diff) medium with the color intensity indicating LabGym’s certainty of that behavior. *n* = 20 spheroids per condition. Right: Percentage of time that spheroids exhibit recruiting behavior over time. *n* = 20 spheroids per condition. **f**, Ranked list of recruiting behavior inhibition following 30 min incubation with compounds that inhibit actin (cytochalasin D, concovalin A), formin (SMIFH2), arp2/3 (CK-666), myosin (blebbistatin), dynamin (dyngo-4a, MiTMAB), Na^+^/H^+^ exchange (DMA, EIPA), PI3K (LY294002), 2^nd^ messenger (U-73122), or MMP (GM6001, marimastat), with EP4 inhibitor L-161,982 used as a positive control in a dose present in differentiation media and Rho kinase inhibitor H-1152 used as a negative control in the dose present in spheroid media. Ratio of inhibition to viability is presented. *n* = 15-18 spheroids per condition. Data are representative of at least 2 independent experiments. Two-tailed Student’s t test in (**e**). One-way analysis of variance (ANOVA) with Tukey’s multiple comparisons post hoc test (**f**). ***p < 0.001, ****p < 0.0001.

The large dynamic range of the recruitment phenotype occurred only in spheroids grown in ISC media (~80% recruiting) unlike those transitioned to differentiation media (~20% recruiting) (Fig. 3e). This platform thus appeared to be suitable for identifying compounds that could inhibit underlying processes to these multicellular behaviors. Bulk transcriptomic datasets suggesting activation of actin filament motor activity may underlie the spheroid behaviors (Extended Data Fig. 5a). We evaluabled compounds that are known to inhibit general actin polymerization/assembly (Cytochalasin D, Concovalin A), formin-dependent nucleation of linear actin filaments (SMIFH2), Arp2/3 complex-dependent nucleation of branched actin filaments (CK-666), myosin II motor (blebbistatin), dynamin (Dyngo-4a, MiTMAB), Na^+^/H^+^ exchange (involved in micropinocytosis and phagocytosis; DMA, EIPA), PI3K (LY294002), 2^nd^ messenger (U-73122), and MMP (GM6001, Marimastat). Several compounds inhibited the recruitment phenotype (Extended Data Fig. 5b) with minimal viability change (Extended Data Fig. 5c), namely those that inhibited actin dynamics including formin and Arp2/3, myosin II, and to a lesser degree, dynamin and MMP (Fig. 3e, Extended Data Fig. 5d for LabGym-quantified data). Since the mechanisms that underlie recruitment are also involved in critical cellular functions such as nutrient absorption and cell division, we deemed that all values below an inhibition-to-viability ratio of 1 as inhibitory (Fig. 3f). In all, the recruitment activity of spheroids is mediated by basolateral membrane rearrangement coordinated by actin, myosin, dynamin, and 2^nd^ messenger interactions. These results suggest that ISCs use multiple mechanical processes to coordinate this behavior.

### Actin-mediated dynamics and C3-opsonization underlie ISC phagocytosis of apoptotic iIECs

Once apoptotic iIECs were within proximity of spheroids, we observed that individual ISCs extended an actin-rich basolateral protrusion that appeared to play a role in enveloping apoptotic iIECs (Fig. 4a). Transmission electron microscopy (TEM) analysis of FACS-purified ISCs including labeled apoptotic iIEC doublets revealed that the ISC produced highly branched basolateral membrane protrusions in the direction of the apoptotic iIEC (Fig. 4b). It is possible that these were outgrowths of protrusions that existed at steady state along the basolateral membrane of ISCs (Extended Data Fig. 6a). We surmised that inhibition of actin dynamics, including actin nucleation, formin-dependent nucleation at barbed ends, and organization into branched networks would each inhibit uptake of apoptotic iIECs; indeed, inhibitors of each of these components of actin dynamics inhibited uptake (Fig. 4c). Notably, pharmacologic inhibition of Na^+^/H^+^ exchange (necessary for macropinocytosis) did not inhibit uptake (Fig. 4c), suggesting that this modality of engulfment was not directly involved in apoptotic iIEC uptake.

**Fig. 4:**
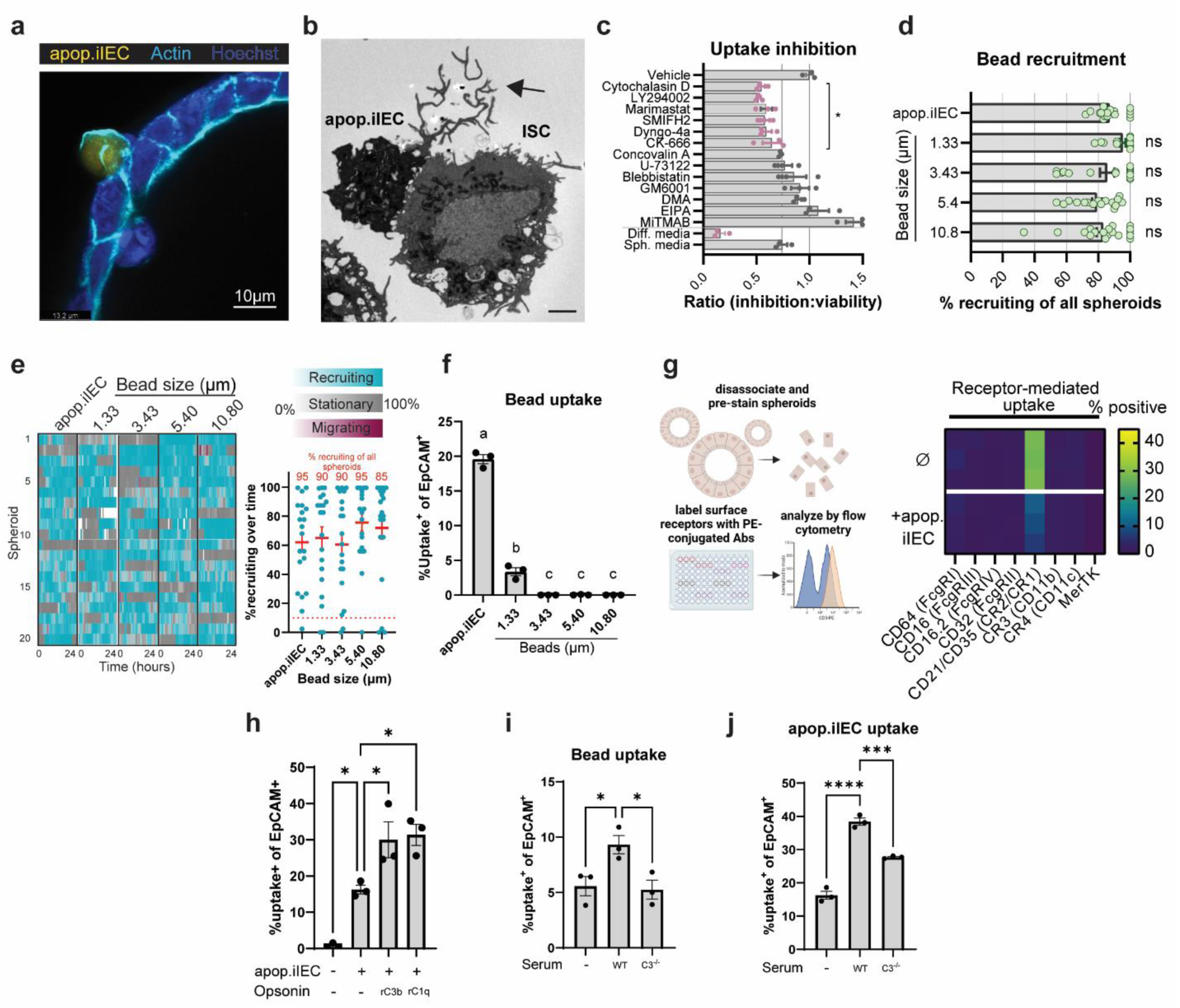
Actin-mediated dynamics and C3-opsonization underlie ISC phagocytosis of apoptotic iIECs. **a**, Merged channels of an immunofluorescently-stained ISC spheroid following 72 h of culture with CellTrace-labeled apop.iIECs. *n* = 3. Scale bar = 10 µm. **b**, Ultrastructure of FACS purified doublet showing ISC (right cell) and apop.iIEC (left cell, appears darker). Arrowhead denotes basolateral membrane protrusion extended by the ISC. Scale bar = 2 µm. **c**, Ranked list of uptake inhibition following 30 m incubation with compounds that inhibit actin (cytochalasin D, concovalin A), formin (SMIFH2), arp2/3 (CK-666), myosin (blebbistatin), dynamin (dyngo-4a, MiTMAB), Na+/H+ exchange (DMA, EIPA), PI3K (LY294002), 2nd messenger (U- 73122), MMP (GM6001, marimastat), with EP4 inhibitor L-161,982 used as a positive control in a dose present in differentiation media and Rho kinase inhibitor H-1152 used as a negative control in the dose present in spheroid media. Uptake was measured by flow cytometry analysis of CellTrace^+^EpCAM^+^ ISCs. *n* = 3 per condition. **d**, Frequency of spheroids that exert recruiting behavior in cultures supplemented with the fluorescently labeled apop.iIECs, and beads of the indicated sizes. *n* = 15-18 per condition. **e**, LabGym quantification of the time that spheroids exhibited Recruiting, Stationary, or Migrating behaviors. (Left) Behavior annotation timeline and (Right) percentage of time that spheroids spent in each behavior, with frequency of recruiting spheroids of all spheroids in red. Red dotted line denotes non-recruiting behavior as determined by spheroids grown in differentiation medium. *n* = 20 spheroids per condition. **f**, Flow cytometric analysis of fluorescent apop.iIEC or bead uptake following single cell, Sytox^-^, and EpCAM^+^ gating. *n* = 3 for each condition. **g**, Cell surface phagocytosis receptor staining with (Left) schematic of experimental pipeline and (Right) expression by percent positive of all ISCs. *n* = 3 for both conditions. **h**, Frequency of uptake^+^ of CellTrace-labeled apop.iIECs that underwent opsonization with recombinant C3b (rC3b) or C1q (rC1q). n = 3 for each condition. **i**, Frequency of uptake^+^ of fluorescent 3.4 µm beads that underwent opsonization with serum from wild-type (WT) or C3^-/-^ mice. n = 3 for each condition. **j**, Frequency of uptake^+^ of CellTrace-labeled apop.iIECs that underwent opsonization with serum from WT or C3^-/-^ mice. n = 3 for each condition. Data are representative of at least 2 independent experiments. One-way analysis of variance (ANOVA) (**c,d,e,f,h,i,j**). *p < 0.05, **p < 0.01, ***p < 0.001, ****p < 0.0001. ns, not significant. Letters denote significant differences among groups in (**f**).

The effect of actin inhibitors on uptake and transcriptomic comparison between uptake^+^ and uptake^-^ cells (Extended Data Fig. 6b) was consistent with multiple methods of cellular uptake. To rule in phagocytosis, defined as the cellular ingestion of particles larger than 0.5 µm through a receptor-mediated process^34^, we tested uptake of inert beads of defined sizes (1-10 µm). Spheroids recruited beads of each size in this range to proximity (see Fig. 3c, Fig 4d). Spheroids could even attempt to recruit beads as large as 120 µm, far larger than what a single cell was capable of engulfing (Supplement video 6). However, uptake of beads was much more limited than that of apoptotic iIECs and appeared to be size-dependent (Fig. 4f). Notably, beads larger than 1 µm were not taken up, including beads that were the size of apoptotic iIECs (~5 µm) (Fig. 4f). This result suggested that ISC recruitment and uptake are governed by different processes. In addition, uptake in the absence of a signal from the engulfed iIEC is limited and size dependent. The bead experiments suggested that a specific receptor(s) initiates uptake, perhaps through phagocytosis. We thus tested the surface expression of common phagocytic receptors using a flow cytometry-based screen (Fig. 4g). Of the receptors tested, the efferocytosis receptor, MerTK, was notably not detected, however there was robust expression of CR1/CR2 at baseline and after apoptotic iIEC supplementation (Fig. 4g). In mice, CR1/CR2 forms a heterodimeric receptor whose ligands include complement factors C3b and C1q. Interestingly, while uptake^+^ ISCs upregulate *C3* transcript (Extended Data Fig. 6b), spheroids generated from *C3*^-/-^ and *C1qa*^-/-^ mice were equally competent in apoptotic iIEC uptake (cultures were supplemented with heat-inactivated FBS) (Extended Data Fig. 6c).

We then tested whether apoptotic iIECs provided complement factor C3. Interestingly, staurosporine treatment for 1 hour resulted in an acute induction of *C3* mRNA (Extended Data Fig. 6d), followed by a robust increase in C3 surface positivity after 24 hours of treatment (apoptotic iIECs) compared to live iIECs (Extended Data Fig. 6e). In addition, C3 opsonization on iIECs appeared stable over several months (Extended Data Fig. 6f). This suggested that apoptotic iIECs produced C3 that promoted self-opsonization during programmed cell death. Indeed, further opsonization with exogenous recombinant C3b and C1q increased their uptake by ISCs (Fig. 4h), whereas incubation of either beads (Fig. 4i) or apoptotic iIECs (Fig. 4j) in serum from *C3*^-/-^ animals did not. Altogether, actin and receptor mediated uptake of apoptotic bodies suggests that the uptake modality is likely phagocytosis.

### Engulfed material in ISCs undergoes intracellular degradation

Considering that engulfed material in other phagocytes is trafficked to and digested in phagolysosomes, we sought to understand whether apoptotic iIEC cargos share the same fate following internalization by ISCs. From TEM of FACS-purified uptake^+^ ISCs, we observed large intracellular vesicles that appeared approximately the same size as the cell’s nucleus (Fig. 5a), a comparison first drawn by Cheng and Leblond^12^. Since internalized material in professional phagocytes undergo lysosomal degradation, we queried this pathway in ISCs. First, internalized fluorescently labeled apoptotic iIECs colocalized with the lysosomal marker LAMP1 (Fig. 5b), with uptake^+^ cells having lower intracellular pH as reported by the pH-sensitive pHrodo dye (Fig. 5c). Second, we found that spheroids degrade internalized exogenous DQ-ova, a substrate that fluoresces upon proteolytic degradation. However, when compared to a macrophage-like cell line, RAW 264.7, DQ-ova degradation was substantially lower at all time points (Fig. 5e). Third, ISCs could diminish fluorescent apoptotic iIEC signal in a span of three days; however, as with DQ-ova, RAW 264.7 cells were far more efficient (approximately 20x) in clearing apoptotic iIECs (Fig. 5f, supplemental video 7; Top: spheroids, Bottom: RAW 264.7 cells). Fourth, uptake^+^ ISCs showed upregulation of genes involved with peptidase and lysosomal acidification (Extended Data Fig. 7a). These results support a model whereby ISCs are capable of degrading phagocytosed material though at a decreased capacity compared to professional phagocytes.

**Fig. 5:**
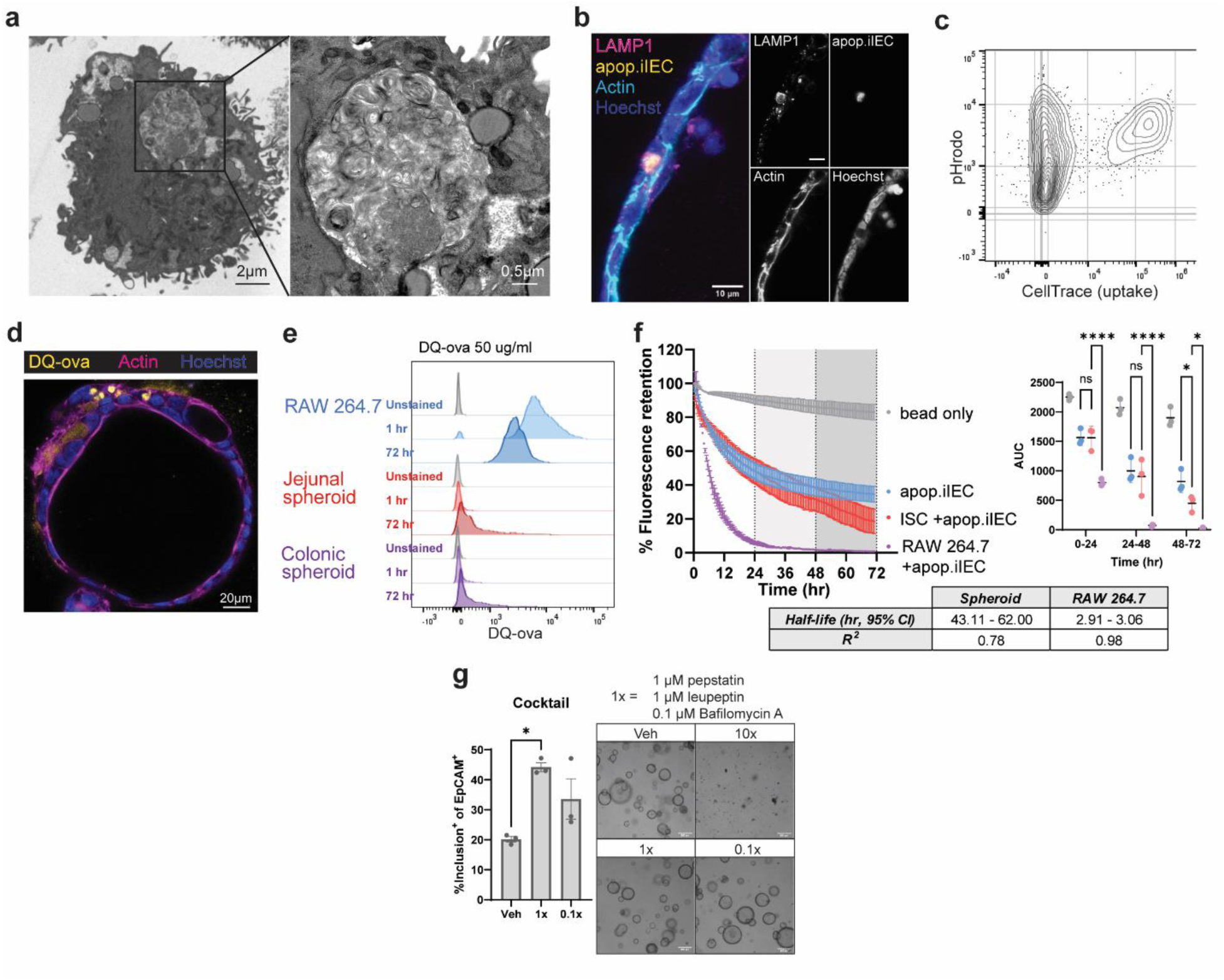
Engulfed material in ISCs undergoes intracellular degradation. **a**, Ultrastructural microscopy of FACS-purified uptake^+^ ISC following 72 h incubation with fluorescently labeled apop.iIEC. Scale bar = 2 µm. (Right) inset of the engulfed material. Scale bar = 0.5 µm. **b**, Immunofluorescence microscopy of a spheroid in culture with apop.iIEC for 72 h. (Left) Merged channels and (Right) individual channels. Scale bar = 10 µm. **c**, Flow cytometry plot of ISCs with uptake plotted against acidification (pHrodo). n = 3 per condition. **d**, Spheroids incubated with DQ-ova for 72 h. Scale bar = 25 µm. *n* = 3. **e**, DQ-ova signal in ISCs or RAW 264.7 cells incubated with DQ-ova for the indicated times. *n* = 3 per condition. **f**, Retention of apop.iIEC fluorescence signal as detected by timelapse fluorescence microscopy. apop.iIECs were incubated with ISCs and RAW 264.7 cells. (Left) Percentage of fluorescence signal compared to time 0. (Right) Area under the curve (AUC) for calculating statistical significance. (Bottom) Half-life of fluorescence signal as a proxy of rate of apop.iIEC degradation by ISCs and RAW 264.7 cells. *n* = 3 for each condition. **g**, Percent of ISCs with apop.iIEC inclusion following incubation of a cocktail of pepstatin, leupeptin, and bafolmycin A at the indicated concentrations. *n* = 3 for condition. Data are representative of at least 2 independent experiments. One-way analysis of variance (ANOVA) with Tukey’s multiple comparisons post hoc test (**f**,**g**). *p < 0.05, ****p < 0.0001. ns, not significant.

We then tested whether degradation in ISCs can be halted by lysosomal inhibitors. Pepstatin, an aspartyl protease inhibitor, and leupeptin, a serine protease inhibitor, both decreased ISC apoptotic iIEC degradation (Extended Data Fig. 7b-c) without affecting spheroid morphology long term. However, both inhibitors elevated intracellular acidification, perhaps in a compensatory manner (Extended Data Fig. 7b-c). Limiting lysosomal acidification using Bafalomycin A1, a vacuolar H^+^-ATPase inhibitor, increased intracellular pH in a dose-dependent manner (Extended Data Fig. 7d). Further, treatment with a cocktail of pepstatin, leupeptin, and Bafalomycin A inhibited apoptotic iIEC degradation by ISCs (Fig. 5g). Altogether, these results indicate that engulfed apoptotic iIECs undergo lysosomal degradation, but that this activity is much slower compared to professional phagocytes.

### ISC phagocytosis of apoptotic bodies enhances CD4^+^ T cell activation

The 2-week q.o.d EdU model leads to crypt-specific epithelial apoptosis is associated with an accumulation of immune cells around crypts (Fig. 1g,h). In addition, EdU enhanced the intestinal lamina propria T_EM_ phenotype (Fig. 1i) and led to production of chemokines that recruit activated T cells by local immune cells (Fig. 1j). One interpretation is that ISCs produce factors that mediate immune enhancement in response to engulfment of apoptotic epithelial cells. Conversely, ISCs may be inefficient at apoptotic cell clearance, leading to secondary necrosis mediated by other cell types. To test the direct involvement of ISCs in modulating immune response, we combined epithelial spheroid culture with an affected cell type, CD4^+^ T cells, via a classic *ex vivo* T cell stimulation assay using Celltrace dilution as a proxy of activation (Extended Data Fig. 8a,b). We observed that supplementing the entire ISC and apoptotic body culture that contains 1) uptake^+^ ISCs, 2) uptake^-^ ISCs, and 3) apoptotic iIECs to freshly purified CD4^+^ T cell culture with αCD3 stimulation yielded suppressed T cell proliferation (Extended Data Fig. 8c). However, the immunomodulatory activity was attributed to apoptotic iIECs (Extended Data Fig. 8d); this response occurred in a dose-dependent manner (Extended Data Fig. 8e). We therefore refined our approach by supplementing FACS-purified uptake^+^ and uptake^-^ jejunal and colonic ISCs (removed non-internalized apoptotic iIECs) and co-cultured each fraction with CD4^+^ T cells (Fig. 6a). Uptake^+^ ISCs enhanced CD4^+^ T cell proliferation compared to uptake^-^ ISCs (Fig. 6a,b) in a dose dependent manner (Fig. 6c). These data show that uptake^+^ ISCs potently enhanced the proliferation of CD4^+^ T cell even at a 1:50 ratio (ISC:CD4^+^ T cells) (Fig. 6c). In addition, uptake^+^ ISCs originating from both mouse colon (Fig. 6d, Extended Data Fig. 9a) and mouse jejunum (Extended Data Fig. 9b) enhanced T cell production of cytokines including those associated with T cell activation (IL-2), Th1 (IFNγ), Th2 (IL-4, IL-5, IL-13), and Treg (IL-10), but not Th17 (IL-17) effector functions (Fig 6d). This response could also be detected by CD4^+^ T cells isolated from mesenteric lymph nodes (Fig. 6e-f), suggesting that T cells from this location can also undergo the same enhanced activity.

**Fig. 6:**
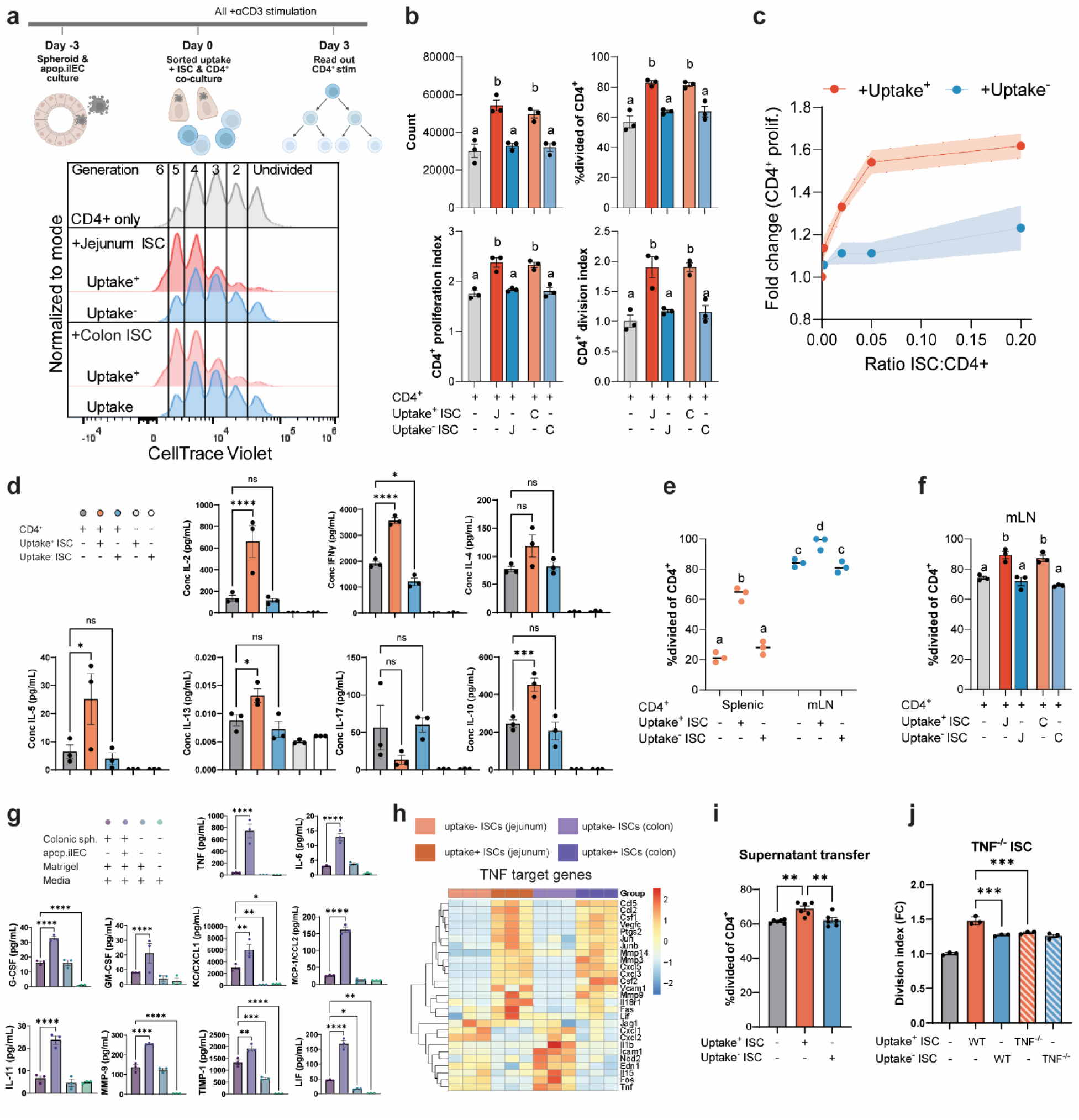
ISC uptake of apoptotic bodies enhances CD4^+^ T cell activity through TNF. **a**,**b**, FACS-purified uptake^+^ and uptake^-^ ISCs from spheroids derived from mouse jejunum or colon tissue were added to freshly isolated and CellTrace-stained splenic CD4^+^ T cells. The two cell populations were co-cultured on aCD3-coated plates for 72 h and underwent flow cytometry analysis with (**a**) proliferation histograms plotted and (**b**) cell proliferation measures calculated from proliferation modeling. *n* = 3 for all conditions. **c**, FACS-purified uptake^+^ and uptake^-^ ISCs were added to freshly isolated and CellTrace-stained splenic CD4^+^ T cells in the indicated ratios with the number of CD4^+^ T cells held constant. The two cell populations were co-cultured on aCD3-coated plates for 72 h and underwent flow cytometry analysis. CD4^+^ proliferation index is plotted with SEM represented as shaded area. *n* = 3 per condition. **d**, Luminex cytokine analysis of ISC and splenic CD4^+^ T cell co-culture supernatant following 72 h of co-culture. **e**, FACS-purified uptake^+^ and uptake^-^ ISCs from spheroids derived from mouse colon were added to freshly isolated and CellTrace-stained CD4^+^ T cells isolated from mouse spleen or mLN. *n* = 3 per condition. **f**, FACS-purified uptake^+^ and uptake^-^ ISCs from spheroids derived from mouse jejunum or colon were added to freshly isolated and CellTrace-stained mLN CD4^+^ T cells. *n* = 3 per condition. **g**, Luminex cytokine analysis of culture supernatant of spheroids were cultured with or without apop.iIEC for 72 h. *n* = 3 for all conditions. **h,** Bulk RNAseq comparing uptake^+^ to uptake^-^ spheroids derived from mouse jejunum or colon. TNF target genes generated from mouse KEGG database. *n* = 3 per condition. **i**, FACS-purified uptake^+^ and uptake^-^ ISCs were cultured in complete RPMI overnight. ISC-conditioned media was collected and supplemented to freshly purified and CellTrace-labelled CD4^+^ T cell culture on αCD3 pre-coated plates. Cell division of assess by flow cytometry. *n* = 3 per condition. **j**, FACS-purified uptake^+^ and uptake^-^ ISCs grown from colon tissue of TNF^-/-^ or WT mice were supplemented to freshly purified and CD4^+^ T cell culture on αCD3 pre-coated plates. *n* = 3 per condition. Data are representative of at least 2 independent experiments. One-way analysis of variance (ANOVA) with Tukey’s multiple comparisons post hoc test (**b**,**c**,**d**,**e**,**f**,**g**,**i**,**j**). *p < 0.05, ***p < 0.001, ****p < 0.0001. ns, not significant. Letters denote significant differences among groups in (**b**,**e**,**f**).

### ISC-produced TNF augments T cell activity

To identify factors that mediated ISC-enhanced immune stimulation, we first compared cell surface expression of specified proteins on ISC-enriched spheroids with or without supplementation of apoptotic iIECs. Of note, we did not observe expression of MHCII (via anti-IA/IE) under either condition. We only observed increased expression of surface molecules with known immune-suppressive roles (e.g. MICL and PDL1) with the addition of apoptotic iIECs (Extended Data Fig. 10a). We next screened expression of a panel of soluble factors in the culture supernatants of spheroids with or without supplementation of apoptotic iIECs. Using a serum-free medium^23^ to reduce the masking of cytokine signals, we found that spheroids cultured with apoptotic iIECs upregulated detectable production of several cytokines including G-CSF, GM-CSF, IL-6, MMP-9, LIF, IL-11, CCL2, TNF, TIMP-1, and MCP-1 (Fig. 6g, Extended Data Fig. 10b).

Of the upregulated cytokines, TNF was notable due to the magnitude of the measured upregulation and its known function in enhancing CD4^+^ T cell activation^35^, which we repeated over a wide dose curve that included the concentration of TNF measured in the serum-free spheroid culture (Extended Data Fig. 10d). Even though *Tnf* transcript was not upregulated in uptake^+^ ISCs suggesting post-transcriptional regulation, many genes associated with TNF signaling were, including chemokines (e.g. *Ccl5* and *Ccl2*), metalloproteases (e.g. *Mmp3*, *Mmp9*, *Mmp14*), and transcription factors (e.g. *Jun* and *Junb*) typically associated with inflammatory processes (Fig. 6h), as were cytokines (Extended data 10c).

We further validated detection of TNF protein using standard stem cell medium containing serum by ELISA (Extended Data Fig. 10e) and found that apoptotic iIECs themselves did not transfer any TNF that may otherwise explain the increase in concentration (Extended Data Fig. 10e). Notably, uptake of apoptotic iIECs by bone marrow-derived macrophages (BMDMs) skewed towards M1 or M2 phenotypes did not elevate TNF or IL-6 secretion (Extended Data Fig. 10f,g), illustrating the functional difference of the response of ISCs to phagocytosis of apoptotic bodies.

To test whether ISC-conditioned media transferred activity to CD4^+^ cells, we incubated FACS-purified uptake^+^ and uptake^-^ spheroids in standard T cell medium and transferred the conditioned media to freshly isolated CD4^+^ T cells. This was sufficient to enhance CD4^+^ T cell activation (Fig. 6i), however the result was modest compared to co-culture, possibly due to the relative short half-lives of cytokines^36^. We next determined the role of epithelial-derived TNF enhancing CD4^+^ proliferation. We cultured intestinal epithelial spheroids from *TNF*^-/-^ mice with apoptotic iIECs and supplemented CD4^+^ T cell culture with either uptake^+^ or uptake^-^ ISCs. We found that ISC-enhanced T cell activity was diminished with uptake^+^ *TNF*^-/-^ spheroids (Fig. 6j). Next, adding neutralizing antibody against TNF to CD4^+^ T cell cultures supplemented with uptake^+^ ISCs was sufficient in reducing the CD4^+^ cytokine response (Extended Data Fig. 10h). These results support a model whereby ISC-produced TNF following phagocytosis mediates enhanced activity of CD4^+^ T cells.

## DISCUSSION

While the word crypt evokes stillness and death, we found that apoptosis within intestinal crypts provokes immune activity. The heart of this process is the ISC, which in addition to its profound role in maintaining the epithelial barrier, is capable of sensing cell death through phagocytosis of apoptotic bodies. Our data suggest that ISCs phagocytose apoptotic bodies, not solely for the purpose of removing debris but also for providing a signal to enhance local immune activity in the intestine. This enhancement orchestrates the local immune system to protect surrounding tissue from damage. This work provides a mechanism that supports an emerging paradigm in the field that ISCs and immune cell populations engage in bidirectional communication to poise immune responses to protect the intestine^37^. Our study underscores the role of ISCs as a sentry in the coordination of physical and immunologic barriers, and the detailed communication between cells of multiple lineages that is required to maintain tissue integrity in the face of ongoing challenges.

A central finding of our study is that ISCs enhance local mucosal immunity through engulfment of apoptotic bodies that are locally produced and include cells derived within the crypt epithelium. This finding stands in contrast to a substantial body of work that supports the model in which efferocytosis is the default mechanism for clearing apoptotic bodies across various tissue niches, a process fundamental to the effector functions of both professional and non-professional phagocytes^29,38^. Efferocytosis is driven in large part by signaling through Tyro3/Axl/MerTK receptors that inhibit a proinflammatory response to apoptotic body uptake^39–41^. MerTK, in particular, mediates downregulation of inflammatory processes, and its deficiency has consequences including inflammation and autoimmune diseases^42,43^. Of relevance to this study, MerTK is critical for the efferocytosis of hair follicle stem cells during the developmentally programmed regression of these structures^14^. Here, we found that ISCs do not express detectable MerTK; these data are further supported by publicly available databases in mouse^44^ and human^45^ and may underlie enhanced immune activity and explain seemingly paradoxical findings previously observed^46^. Moreover, ISCs exhibit properties that distinguish them from other non-professional phagocytes. Notably, ISCs are located in crypts that typically do not experience waves of programmed cell death; cell death in these areas occurs primarily in response to specific triggers such as injury, mutation, or infection^47^. Also, in contrast to intestinal crypts, differentiated intestinal villus cells frequently undergo programmed cell death, which is often followed by their extrusion into the lumen as well as uptake by local macrophages, the latter resulting in efferocytosis^20^. Furthermore, apoptosis on the villus (but not crypts) can be ramped up by treatment with poly(I:C) without triggering a pro-inflammatory response^48^. Therefore, the presence of excess apoptotic bodies in the crypt is unusual. The model that emerges from our data is that ISCs sense cell death in the crypt through engulfment of apoptotic bodies, which in turn rapidly sends a signal for multi-modal fortification of the barrier in this location. The balancing act the intestine is to maintain an appropriate level of host-mediated immune activity over time in a predominantly tolerogenic tissue. Conversely, this model reveals a potential source of pathology: excessive apoptosis in the crypt may contribute to diseases characterized by hypertrophy of the intestinal epithelium, ultimately exacerbating local immune responses and fueling chronic inflammation within the tissue^49^.

Our findings suggest that ISCs possess multiple cell-intrinsic properties that support their capabilities to be nonprofessional phagocytes. Through 3D intestinal spheroid cultures, we discovered that ISC spheroids can migrate as a cohesive unit directly toward apoptotic iIECs, shedding light on the mechanisms driving ISC phagocytosis of apoptotic cells. Directional migration has been observed in other mammalian epithelial organoid systems^50,51^ and is a property that may originate from developmental programs that play a role in processes such as wound healing^52^. We found that the spheroids moved through Matrigel at a speed of ~20 µm per hour, which approximates a similarly rapid rate at which epithelial cells migrate up the crypt-villus axis^53,54^; as well as a rate reported in epithelial wound repair^55^. Further, this supports the dynamic nature of crypts as observed in multi-photon experiments of the live intestine^56,57^. The elegant pulse-chase experiments that Cheng and Leblond published 50 years ago permitted observations of apoptotic body remnants inside ISC progeny^12^. Our data indicate that ISC degradation of apoptotic bodies is approximately 20-fold slower than that of professional phagocytes. This finding, in combination with the data that immune cell numbers are enhanced in the villus core, suggests that ISC phagocytosis of apoptotic bodies in the intestinal crypts can be relayed to the villus to enhance immune cells in this location. We also observed that non-migratory ISCs exert a near-constant vacuum that can draw inert material to proximity of spheroids. We found that spheroids recruited inert apoptotic iIECs and even nonbiologic inert beads from over 100 µm away. It is possible that this vacuuming activity is driven by spheroids pulling on Matrigel fibers via actomyosin-mediated cortical flow. The vacuuming ability, together with their ubiquitous location in every crypt along the intestinal tract and proximity to apoptotic bodies in the crypt, suggest the competitive advantage that ISCs possess in engulfing apoptotic bodies in this tissue niche.

Utilizing a classical CD4^+^ T cell activation assay, we found TNF is a mediator of uptake^+^ ISC-driven immune enhancement. We detected ISC-derived TNF protein through multiple methods and demonstrated functional activity in both *in vitro* and *in vivo* systems. To detect this TNF signal, which can be masked by standard ISC growth medium, we used freshly prepared, serum-free spheroid culture medium. It has been proposed that IECs engage in short range communication and produce lower amounts of cytokine as compared to traditional APC/T cell interactions in culture to enhance immune responses. Further, detecting TNF at the transcript level can be complicated by posttranscriptional regulation including cleavage of its transmembrane form to produce soluble TNF^58^. Nevertheless, several lines of evidence that epithelial cells produce TNF include: 1) *Tnf* transcript were detected in cells at the crypt base^59^, 2) IECs are the major source of TNF in the ileum of TNF^ΔARE^ mice that are susceptible to Crohn’s-like ileitis^60^, and 3) targeted ablation of TNF in intestinal epithelial cells using *Villin*^CreERT2^*; Tnf*^F/F^ mice demonstrated that IEC-derived TNF regulates mucus-producing cells^61^. Our work suggests that in addition to the canonical role of TNF in inducing cell death^62^, IEC production of TNF mediates immune communication and enhances T cell activation^35^ and immune system readiness in the intestinal lamina propria^63^. This implies that while αTNF therapies (i.e. adalimumab, infliximab, etanercept, etc) have been used successfully in the clinic for over 50 years, roles of epithelium-derived TNF are poorly understood and require more specialized and enhanced detection tools.

Although the focus of this work was on the cells that engulf apoptotic bodies, we found that iIECs produce C3 during the apoptosis process, perhaps to influence their own fate and accelerate clearance after life. To our knowledge, this is the first time this was observed. Supporting this, IEC production of C3 has been corroborated by other groups, including in the human Caco-2 IEC line^64,65^, primary organoid culture^66^, and in the villus tips of mice^67^. Consistently, upregulated *C3* transcripts were observed in inflamed mucosal areas of Crohn’s Disease patients^68^, in murine intestinal injury models of DSS^69^ irradiation and ischemia/reperfusion^66^, and in a NF-κB-dependent manner downstream of TNF in a mouse model of LPS-induced sterile inflammation^4^. Upregulated production of C3 in these models of excess cell death in the intestine supports the notion that IECs produce C3 under stress and during programmed cell death. Downstream, C3 opsonization of dead cells can regulate their clearance^70^ and presentation to CD4^+^ T cells^71^; consequences of C3 deficiency include gastrointestinal phenotypes including increased epithelial permeability^67^.

This work debuts a mouse model of intestinal crypt apoptosis induced by repeated administration to the thymidine analog EdU, a common lineage tracing and proliferation reagent. Previous studies showed the genotoxicity of EdU at experimentally-used doses that was more potent than irradiation^72^ and other thymidine analogs including BrdU^73^. The EdU model fulfilled our criteria for developing an *in vivo* model of crypt-localized apoptosis: 1) intestinal epithelial cell apoptosis localized to crypts, 2) non-immune initiation of epithelial cell death, 3) persistence of a healthy population of ISCs, and 4) facile labeling of partially degraded apoptotic bodies. These criteria ruled out many commonly utilized intestinal injury models such as DSS, trinitrobenzene sulfonic acid (TNBS), and Poly(I:C) for widespread and poor localization of cell death; *IL-10*^-/-^ or *TNF*^ΔARE^ for harboring immune compartment perturbations; genetic models targeting ISCs; irradiation and chemotherapeutics for difficulty in labeling dead cells; other thymidine analogs such as BrdU and IdU due to lower genotoxicity and harsh sample procedures necessary for detection. This model sheds light on a potentially unintended side effect of using this common lineage-tracing reagent but also presents opportunities to rigorously study epithelial regeneration following targeted cell death.

Organized immune protection is known to occur in 8-10 Peyer’s patches and hundreds of inducible lymphoid follicles in an adult mouse, calling into question of how these few sentries can protect the length of the gut. Our work shows that ISCs actively participate in their own protection at a local level away from secondary and tertiary lymphoid structures. This local control is an alarm bell existing in every crypt along the intestinal tract (approx. 1 x 10^6^ crypts in adult mouse) that can precede barrier breach. Apoptotic cell phagocytosis serves as an apex signal that leads to ISC-mediated enhanced local immune activity.

## METHODS

### Mice

C57Bl/6J (Jackson Labs, Stock #000664), B6.129S4-C3tm1Crr/J (C3^-/-^, Jackson Labs, Stock #029661), and B6.129S-Tnftm1Gkl/J (TNF^-/-^, Jackson Labs, Stock #005540) mice were purchased from Jackson Laboratory. Tissue from C1q^-/-^ mice was a kind gift from Dr. Feng Lin (Cleveland Clinic Lerner Research Institute). All mice were on the C57Bl/6 background and age and sex-matched male and female mice 8-15 weeks of age were used to perform experiments. All experiments involving mice were done with littermate controls. Mice were housed in a 11 h light/13 h dark cycle specific pathogen-free facility and fed standard chow *ad libitum*. Enrichment and privacy were provided in all cages with nestlets and an autoclaved hut. All animal studies and housing were performed in accordance with the guidelines established by the Institutional Animal Care and Use Committee at Cleveland Clinic Lerner Research Institute.

### EdU model of intestinal crypt apoptosis

Age and sex-matched mice between 8 and 15 weeks were treated q.o.d with EdU (Sigma). EdU was resuspended in DMSO at a concentration of 50 mg/ml and administered to mice at a dose of 100 mg/kg diluted to 1 ml sterile PBS via i.p. injection. Vehicle controls were administered in an equivalent volume of DMSO according to the same schedule. Mice were weighed and fecal samples were collected at every treatment time point, which occurred within a 2 h window from the initial start on day 0. Mice were sacrificed at various time points during EdU treatment (24 h, 7 d, 14 d). For the 14 d timeframe, one additional dose of EdU was administered 24 h prior to harvest. At the conclusion of the studies, mice were euthanized, blood samples were collected via cardiac puncture, and serum was obtained through centrifugation at 10,000 xg at 4°C for 10 m. Spleens, mesenteric lymph nodes, and small intestinal Peyer’s patches were collected for downstream analyses. Intestinal lamina propria mononuclear cells were isolated by incubating 20 cm lengths of flushed and filleted intestinal samples without Peyer’s patches in EDTA (Gibco) followed by Collagenase type VIII (Sigma). Intestinal samples with Peyer’s patches intact were fixed in 4% paraformaldehyde (PFA) for histologic processing. The EdU-Click-it kit (Thermofisher) was used to detect EdU signal.

### Barrier integrity

Intestinal barrier integrity was assessed by FITC-dextran as described previously^48^. Briefly, mice were fasted mice for 4 h. 4-kDa FITC-dextran (Sigma) was dissolved in PBS to make 100 mg/mL, and 44mg/100g was delivered by oral gavage. 3 h after gavage, blood was collected by cardiac puncture, and serum was obtained following centrifugation. FITC-dextran levels were measured (ex: 485 nm/em: 528nm) based on a standard curve on an Biotek Cytation 5 (Agilent) microplate reader. Treatment with Poly(I:C) via i.p. injection followed by harvest after 6 h was used as a positive control for small intestinal permeability^48^. 1mg/mL poly(I:C) HMW (InvivoGen) was resuspended according to the manufacturer’s instructions and 20 mg/kg was administered via i.p. injection.

### Mouse intestinal spheroid lines

Mouse colonic stem cell spheroid lines were derived from 8 to 15-week-old mice with indicated genotypes as previously described^21^. Briefly, a 1 cm segment of intestinal tissue was minced and digested in 2 mg/ml Collagenase I solution (Invitrogen) with Gentamicin (Sigma) and incubated at 37°C for 20-30 min, with vigorous pipetting every 5 m. The crypt compartment was collected by centrifugation, washed twice with DMEM/F12 media (Sigma), resuspended in cold Matrigel (Corning) and plated onto a 24-well tissue culture plate and cultured in 50% L-WRN conditioned medium supplemented with 10uM Rock inhibitor Y-27632 (R&D Systems).

### Generation of apoptotic immortalized mouse intestinal epithelial cells

To generate immortalized cell lines, we transfected HEK293T cells with plasmid data from pRRLsin-SV40 T antigen-IRES-mCherry (Addgene #58993), packaging plasmid (Addgene #12260) and envelope plasmid (Addgene #12259) with Jetprime transfection reagent (Polyplus). Virus was collected and concentrated using PEG-it (System Biosciences) centrifugation precipitation and was used to infect spheroids. Spheroids were allowed to recover in Matrigel and 50% L-WRN media. Following expansion of transfected cells, spheroids were adapted to grow on plastic and in DMEM media. Apoptotic cells were generated by supplementing cell culture with 10 µM staurosporine (Cayman) to induce apoptosis^74^ and 1 unit/mL DNAse I (New England Biolabs) to prevent cells from clumping. To fluorescently label apop.iIECs, cells were washed in PBS and stained with CellTrace Violet or CellTrace Green (ThermoFisher) according to the manufacturer’s recommendations. Cells or beads (Spherotech) were opsonized with 20% mouse serum or 1mg/ml recombinant C3 or C1q in GVB^++^ buffer containing Ca^++^ and Mg^++^ (Complement Technology) for 37°C for 30 m, then quenched with 10% by volume 100 mM EDTA.

### Spheroid and ALI culture

Adult mouse intestinal spheroids were maintained by passaging every 3-4 d. To passage, spheroids were disassociated using TrypLE (Gibco) along with vigorous pipetting and plated in cold Matrigel. Stem cell media contained 50% L-WRN conditioned medium, which is a 1:1 mix of L-WRN conditioned medium (CM) and fresh primary culture media, which is Advanced DMEM/F- 12 (Invitrogen) supplemented with 20% fetal bovine serum, 2 mM l-glutamine, 100 units/ml penicillin, and 0.1 mg/ml streptomycin supplemented with 10 μM Y-27632 (ROCK inhibitor; R&D Systems). Enterocyte differentiation medium^22^ was primary culture medium without serum supplemented with 50 ng/ml EGF (Peprotech), 10 μM Y-27632 (ROCK inhibitor; R&D Systems), and 10 μM L-161,982 (EP4 inhibitor, R&D Systems). To induce synchronized differentiation, stem cell-enriched spheroids were grown in spheroid media for 48 h and then transitioned to differentiation media. Spheroids were supplemented with apoptotic cells of fluorescent beads of defined sizes (Spherotech) mixed into the Matrigel. Defined serum-free W/A media ^23^ was supplemented with R-spondin (500 ng/ml), Noggin (100 ng/ml), and 1:100 HEPES. Unless otherwise noted, apoptotic cells were imIEC cell lines added at the time of splitting at a 1:1 ratio. Cytochalasin D (5 µM), Concovalin A (10 µM), SMIFH2 (10 µM), CK-666 (10 µM), Blebbistatin (10 µM), Dyngo-4a (30 µM), MiTMAB (30 µM), DMA (100 µM), EIPA (100 µM), LY294002 (40 µM), U-73122 (50 µM), GM6001 (50 µM), and Marimastat (10 µM) were individually applied to spheroid culture for 30 m prior to culture. DQ-ova (50 µg/ml), leupeptin (10 µM), pepstatin (1 µM), and bafilomycin A (100 nM) were individually provided for the duration of the culture. LysoTracker (50 nM) was added to the last hour of culture prior to analysis.

ALI culture was performed as previously reported^31^. Briefly, spheroids were dissociated into single cells using TrypLE Express (ThermoFisher) digestion and seeded onto Transwells (Corning) pre-coated by 10% Matrigel (Corning) for 20 m. ~20,000 cells were seeded per well. Initially, cells were submerged in 150 μL 50% L-WRN medium with 10 uM Rock inhibitor Y-27632 (R&D Systems). Medium in the upper chamber of the Transwell was removed after 7 d to create an air-liquid interface (ALI). Cells were grown in ALI with 50% L-WRN medium in the bottom chamber for additional 7 d with medium changes every 3 d.

### Primary immune cell purification

Primary cells were isolated from freshly harvested mouse spleens or mesenteric lymph nodes and reduced to a single cell suspension by passing them through a sterile 100 µM nylon mesh cell strainer (Fisher Scientific). The single cell suspensions were labeled with anti-mouse CD4 magnetic microbeads (Miltenyi Biotec) and separated by magnetic column according to the manufacturer’s recommendations. Primary neutrophils were isolated from freshly drawn blood. After centrifugation and RBC lysis (BD), cells were labeled with anti-mouse Ly6G (Miltenyi Biotec) and separated by magnetic column.

### Cell culture

After magnetic sorting, CD4^+^ T cells were cultured in 96-well round bottom plates (Corning) at 50,000 cells per well in advanced RPMI (Gibco) supplemented with 10% heat-inactivated fetal bovine serum, 55 µM β-mercaptoethanol, 100 U/mL and 100 µg/mL Penicillin/Streptomycin, and 0.2 mM L-glut. Plates were pre-coated with αCD3ε (eBioscience) 2.5 µg/mL diluted in PBS and then incubated at 4°C overnight followed by two washes with PBS before receiving cells. Recombinant mouse TNF (R&D Systems) was added to culture at the indicated concentrations.

To grow BMDMs, bone marrow was harvested from mouse femurs and plated on petri dishes with RPMI supplemented with 10% fetal bovine serum, L-glut, and penicillin/streptomycin, as well as 25% L929-supplemented media containing macrophage-differentiating growth factors. Additional media with 25% L929 media was supplemented after 3 d of culture, and the media was changed completely on day 6. 7 d of differentiation after initial plating, petri dishes were washed, and adherent macrophages were detached and used in experiments.

iIEC, RAW 264.7 (ATCC TIB-71), L929 (ATCC CCL-1), J774A.1 (ATCC TIB-67), were grown in DMEM supplemented with 10% FBS, L-glut and Penicillin/Streptomycin. B16-F10 (ATCC CRL-6475) cells were grown in DMEM 10% FBS, 1:100 Pen/strep, 1:100 L-glut, and 1:100 sodium pyruvate. CT26 (ATCC CRL-2638) cells were grown in RPMI supplemented with 10% FBS, 1:100 HEPES, 1:100 Pen/Strep, 1:100 L-glut, 1:100 BME, 1:100 NEAA, and 1:100 Sodium pyruvate. All cells were grown in 5% CO2 and 37°C.

### Uptake^+^ ISC and CD4^+^ T cell co-culture

Spheroids were cultured with CellTrace-labeled apop.iIECs for 3 d, after which spheroids were harvested with Cell Recovery Solution (Corning) and digested into a single cell suspension with TrypLE Express (ThermoFisher). Cells underwent FACS sorting (methods below) into uptake^+^ (single cell, EpCAM^+^, CellTrace^+^) or uptake^-^ (single cell, EpCAM^+^, CellTrace^-^) fractions. Sorted ISCs were centrifuged at 300 xg for 5 m and incubated with freshly harvested and purified CD4^+^ T cells from the spleen or mLN of mice. 50,000 CellTrace stained CD4^+^ T cells (in a different CellTrace fluor used to stain apop.iIECs) were seeded per well of a round-bottom 96-well plate, and ISCs were supplemented at a default of 5,000 ISCs to 50,000 CD4^+^ T cells (1:10 ratio) or in the ratios as indicated. Cells were cultured in CD4^+^ T cell culturing conditions. Purified anti-mouse TNF (Biolegend) or Rat IgG1, κ isotype control (Biolegend) were used at a concentration of 2.5 µg/ml for the duration of the co-culture. Cells underwent flow cytometric analysis and supernatants underwent cytokine analysis.

### Flow cytometry

To obtain single-cell suspensions of mouse intestinal lamina propria tissue for flow cytometry analysis, the distal half of the small intestine was excised, and contents were flushed with PBS. Peyer’s patches were carefully excised using fine-tip curved scissors and intestines were filleted longitudinally and cut into approximately 1 cm pieces. Tissue was incubated in pre-warmed 5 mM EDTA in PBS at 37°C shaking at 180 rpm for 15 m. After aspiration of the solution, the previous incubation was repeated. This tissue was minced, transferred into a fresh tube and washed with complete RPMI and allowed to sit for 5 m to inactivate excess EDTA. The tissue was then transferred to digestion media containing collagense type VIII (Sigma) and Dispase (BD) and incubated in a shaking incubator in a horizontal position at 180 rpm for 30 m at 37°C. Tissue was passed through a 70 µm strainer and washed with 5mM EDTA in PBS. The remaining tissue was incubated again with digestion media and added to cells from the first pass through. Cells were spun at 300 xg for 10 m and stained. To obtain single-cell suspensions of Peyer’s patch, mLN, and spleen, tissues were harvested and passed through a 70 µm strainer. Spleen cells underwent RBC-lysis (BD).

Cells underwent Fc blockade using TruStain FcX (Biolegend). Commercial antibodies used included: CD45 – FITC, CD8 – PE, CD44 – PE-Cy7, CD4 – APC, CD3 – APC-Cy7, CD62L – BV711, F4/80 – FITC, B220 – PE, Ly6G – PE-Cy7, CD11c – APC, CD11b – APC-Cy7, IA/IE – BV421, CD45 BV711, EpCAM – PE-Cy7, CD4 – PE, C3 – FITC, EpCAM – FITC, CD64 (FcγRI) – PE, CD16 – PE, CD16.2 (FcγRIV) – PE, CD32 (FcγRII) – PE, CD21/CD35 (CR2/CR1) – PE, CD11b (CR3) – PE, CD11c (CR4) – PE, MerTK – PE, IFNAR-1 – PE, IFN-γR β chain – PE, CD25 (IL-2R) – PE, CD124 (IL-4Rα) – PE, CD132 (common γ chain) – PE, CD126 (IL-6Rα chain) – PE, CD210 (IL-10 R) – PE, IL-33Rα (IL1RL1, ST2) – PE, CD120a (TNF R Type I/p55) – PE, CD282 (TLR2) – PE, CD284 (TLR4) – PE, CD285 (TLR5) – PE, CD40 – PE, CD70 – PE, PDL1 – PE, PDL2 – PE, GITRL – PE, CD80 – PE, CD86 – PE, CD206 – PE, Dectin1 – PE, MICL – PE, CD1d – PE, MR1 – PE, CD115 – PE, Notch1 – PE, Notch2 – PE, Notch3 – PE, Notch4 – PE, Tim4 – PE, OX40L – PE, HVEM – PE, CD153 – PE, CD44 – PE, IA/IE – PE, CD31 – PE, Siglec F – PE, P-Selectin – PE, NK.1 – PE, Rat IgG2a, κ – PE, Armenian Hamster IgG – PE, Rat IgG2b κ – PE, Mouse IgG1, κ – PE, Rat IgG1, κ – PE. Additional flow cytometry reagents include: Sytox Red (Invitrogen), Sytox Green (Invitrogen), Sytox Blue (Invitrogen), Fixable Viability Stain 510 (BD), CellTrace Violet (Invitrogen), CellTrace Green (Invitrogen), Ultracomp compensation eBeads (BD), and Arc/Amine reactive compensation beads (BD).

Cells were analyzed using an Attune NxT flow cytometer (ThermoFisher). Cells were sorted using a FACSAria Fusion (BD) or Bigfoot (ThermoFisher). Analysis of all samples was completed in FlowJo v10.

### Cytokine analysis ELISA and Luminex, Legendplex

Cytokine concentrations were analyzed by LegendMAX sandwich ELISAs performed according to manufacturer’s instructions (Biolegend). Signal was detected with a Biotek Cytation 5 plate reader (Agilent). For Legendplex assays, cytokines analyzed by Legendplex kits (Biolegend) according to manufacturer’s instructions (Biolegend) and read on an Attune NxT flow cytometer. For Luminex assays, media were snap frozen in liquid nitrogen and submitted to Eve Technologies (Calgary, AB) for mouse 44-plex and mouse cardiovascular disease panel cytokine analysis.

### Tissue Preparation and Histology

Dissected intestinal tissue cut to lengths of 10 cm were pinned out and fixed in 4% paraformaldehyde (PFA) overnight at 4°C. The fixed tissue was washed in PBS 3 times before dehydration in 70% ethanol. Tissues were embedded in 2% agar and submitted for paraffin embedding, sectioning, and hematoxylin and eosin staining to the Cleveland Clinic Imaging and Microscopy core. ALI cultured cells were fixed in 4% PFA for 30 m at room temperature and treated similarly to the tissues for histological examinations.

### Immunostaining

Slides underwent de-paraffinized, rehydration, and heat-induced antigen retrieval in Tris-EDTA Buffer pH 9.0 (Abcam) for 20 minutes. Slides were blocked in PBS containing 5% serum and 0.1% Triton-X for 1h at room temperature, and then treated with primary antibodies at desired dilutions for overnight at 4°C. Slides were then washed 3 times in PBS. For immunofluorescence, slides were incubated with secondary antibodies conjugated with fluorophores (ThermoFisher) for 1 h at room temperature. Counterstains used included Hoechst 33342 (Sigma), RedDot (Biotium), and phalloidin conjugated to AF488 of AF647(ThermoFisher). For whole mount staining of spheroids, cells were plated in Matrigel on coverglass culture dishes (Ibidi) and fixed in 4% PFA for 30 min at room temperature. Cells were then blocked, incubated with primary antibodies and followed the same procedures used on slide sections. For immunohistochemistry, VECTASTAIN Elite ABC HRP Kit and I DAB Peroxidase (HRP) Substrate Kit (Vector Laboratories) were used to develop signal. Slides were counterstained with dilute Gill 2 hematoxylin (ThermoFisher) and dipped in bluing reagent (Abcam).

### Epifluorescence and confocal microscopy

Bright field histological images and images of spheroid culture were acquired with a Zeiss Axio Observer 7 inverted fluorescence microscope. Fluorescently labeled slides were imaged with an inverted Leica SP8 confocal microscope using a 40x oil immersion objective. Whole mount spheroids were imaged using an inverted Leica SP8 confocal microscope adapted with a Leica 40x water immersion lens that allows for a high working distance. Z stacks were imaged with system-optimized steps and 3D reconstruction was completed in Leica LAS X 3D software. FIJI (Fiji is just ImageJ) software was used to uniformly adjust brightness and contrast, recolor images, add scale bars and time stamps, and quantify fluorescent intensity.

### Electron microscopy

Sorted ISCs were washed once in PBS and submitted to the Imaging and Microscopy core at CCF for professing. Samples were fixed overnight at 4°C in 2.5% glutaraldehyde + 4% paraformaldehyde in 0.2 M sodium cacodylate buffer (pH 7.4). Fixative was removed and samples were washed with sodium cacodylate buffer. Samples were incubated with 1% osmium tetroxide in H2O for 1 hr at 4°C, washed with sodium cacodylate, then followed by a 1 hr incubation in 1% uranyl acetate in maleate buffer and washing with maleate buffer. Samples were dehydrated and embedded in pure eponate 12 medium. Ultra-thin sections (85 nm) were cut with a diamond knife and stained with uranyl acetate and lead citrate. Sections were observed with a Tecnai G2 Spirit BioTWIN transmission electron microscope operated at 80 kV. Images were captured using a GATAN RIO16 camera.

### Time-lapse microscopy and machine learning-based spheroid behavior analysis by LabGym

Cells were imaged on a Zeiss Axio Observer 7 inverted fluorescence microscope adapted with environmental controls including a motorized heated stage and CO_2_ module. Images were captured every 30 m for up to 72 h and were exported as native .czi files for processing. In FIJI, channels were appropriately assigned, brightness and contrast were uniformly adjusted, and scale bars and time stamps were appended per frame. Images were exported as .tiff files and videos were exported as .avi files for further processing. Quantification of videos was performed using a pipeline previously reported^32,33^. Briefly, spheroids in images were annotated using Roboflow^75^ (2,391 images that were augmented by random rotation and exposure) and used to train a LabGym Detector to detect spheroids in videos (8,000 iterations, inferencing frame size 1,920 pixels). Examples of Recruiting, Migrating, and Stationary behaviors were generated from videos using the trained LabGym Detector, and were manually categorized to train a LabGym Categorizer (complexity set to 3, input size 1,920 pixels). The trained LabGym Detector and Categorizer was then used to automatically analyze spheroid behavior and generate heatmaps.

### RNA isolation and quantitative PCR

RNA was extracted from FACS purified ISCs using the RNeasy Micro Kit according to manufacturer’s directions. Briefly, cells were centrifuged and lysed in RLT buffer with 2- mercaptoethanol. RNA was extracted and eluted in 14 ul of nuclease free water and stored at −80°C. RNA was quantified on a Cytation 5 by diluting 1:1 with nuclease free water. cDNA was prepared using 1ug of starting RNA and using iScript reverse transcriptase kit (Bio-Rad) according to manufacturer’s directions. Quantitative PCR reaction was conducted with TB Green Advantage qPCR 2X- premix (Takara) using a Bio-Rad CFX Connect Real-Time PCR Detection System. Primers were validated for specificity using a water control and conducting melt curve analysis. Data were analyzed using the ΔΔCt method. Primer sequences for quantitative PCR used in this study are as follows; B2m (forward): ACAGTTCCACCCGCCTCACATT, B2m (reverse): TAGAAAGACCAGTCCTTGCTGAAG, C3 (forward): CGCAACGAACAGGTGGAGATCA, C3 (reverse): CTGGAAGTAGCGATTCTTGGCG.

### Bulk RNA sequencing and analysis

RNA was extracted as described above. RNA quality was assessed by Qubit and Tapestation. Library prep was done with SMARTer® Stranded Total RNA-Seq Kit v3 - Pico Input Mammalian and 50 million reads were obtained per sample. Sequencing datasets were produced using an Illumina NovaSeq 6000 instrument. For analysis, reads were aligned to the reference mouse genome (GRCm39) using STAR software^76^. Gene counts were derived from uniquely aligned reads. To identify differentially expressed genes, DESeq2’s median of ratios was used to normalize for sequencing depth and for calculation of fold changes and p-values (nominal & adjusted)^77^. RNA sequencing data will be deposited upon manuscript acceptance.

### Statistics and reproducibility

Statistical parameters including the exact value of n, precision measures (mean ± SEM) and statistical method and significance are reported in the Results and Figure Legends. Data are judged to be statistically significant when p < 0.05. In figures, asterisks denote statistical significance: *, p < 0.05; **, p < 0.01; ***, p < 0.001; ****, p < 0.0001. In some graphs, letters were used to denote significant differences among groups.

## ACKNOWLEDGEMENTS

We thank R. Musich and A. Job (Bioinformatics, CCF Department of Inflammation and Immunity), and M. Yin (Electron Microscopy Core, CCF), M. Lim (Genomics Core, CCF), and S. Rodgers and E. Schultz (Flow Cytometry Core, CCF) for their technical support. This work was supported by the following: NIH (F32DK136180 to J.Y.Z. and R01CA257523 to T.S.S. and O.H.Y.).

## AUTHOR CONTRIBUTIONS

J.Y.Z. and T.S.S. conceptualized and supervised the study. J.Y.Z. designed, performed, and analyzed the data for most experiments. Q.L. designed and created the SV40-immortalized mouse intestinal epithelial cell lines. Y.Hu assisted with LabGym installation and analysis. S.F. assisted with W/A intestinal spheroid culture platform. S.T.E. assisted with epifluorescence microscopy instrumentation. M.J.E., K.J.L., P.E.K., and Y.Han assisted with mouse experiment harvests. H.S. and O.H.Y. contributed expertise on lysosomes and critical discussion. R.E.S. helped with interpretation of TEM images. D.J.S. helped with EdU experiment design and mouse harvests. A.I.I. contributed expertise on cytoskeleton biology. J.Y.Z. and T.S.S. wrote the manuscript. All authors contributed to the critical review of the manuscript.

## EXTENDED DATA FIGURES

**Extended Data Fig. 1 (corresponds to Fig. 1):**
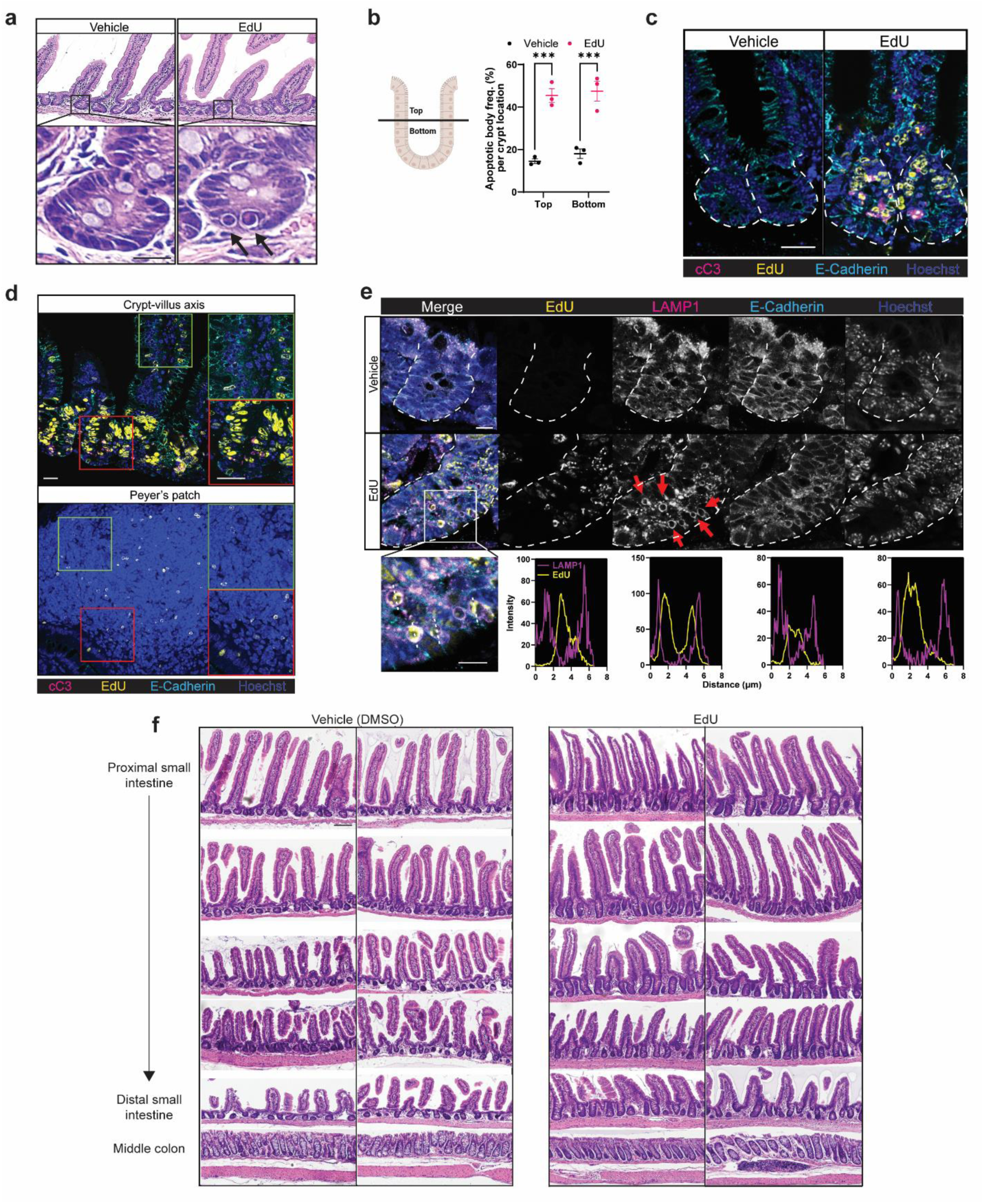
EdU exposure as a model of excess apoptosis in the intestinal crypts. **a,** Representative images of H&E-stained sections of proximal small intestine treated or untreated with 24h i.p. injection of EdU. *n* = 3 (vehicle), *n* = 5 (EdU). Top panels: Scale bar = 50 µm. Bottom panels (insets): crypts with arrows denoting apoptotic bodies. Scale bar = 10 µm. *n* = 3-5 mice per treatment group. **b**, Top: schematic of apoptotic body quantification strategy. Bottom: frequency of crypts containing apoptotic bodies per location (top or bottom of crypt). *n* = 100-200 crypts evaluated per mouse, 3-5 mice per treatment group. **c**,**d**,**e**,**f**, Mice were injected q.o.d with EdU for 7 days. *n* = 3-5 mice per treatment group. (**c**) Representative sections of proximal small intestine and (**d**) with (Peyer’s patch) stained for the indicated targets. Scale bars = 25 µm. (**e**) Representative sections of proximal small intestine with white dotted lines denoting crypts. Scale bar = 50 µm. Magnification of left inset with white dotted line indicating intensity of LAMP1 and EdU calculations. Scale bar = 10 µm. **f**, Representative H&E histology along the length of the mouse intestine from proximal (top) to distal (bottom) (scale bar = 10 µm). n = 3-5 mice per treatment group. Data are representative of at least 2 independent experiments. ***p < 0.001.

**Extended Data Fig. 2 (corresponds to Fig. 1):**
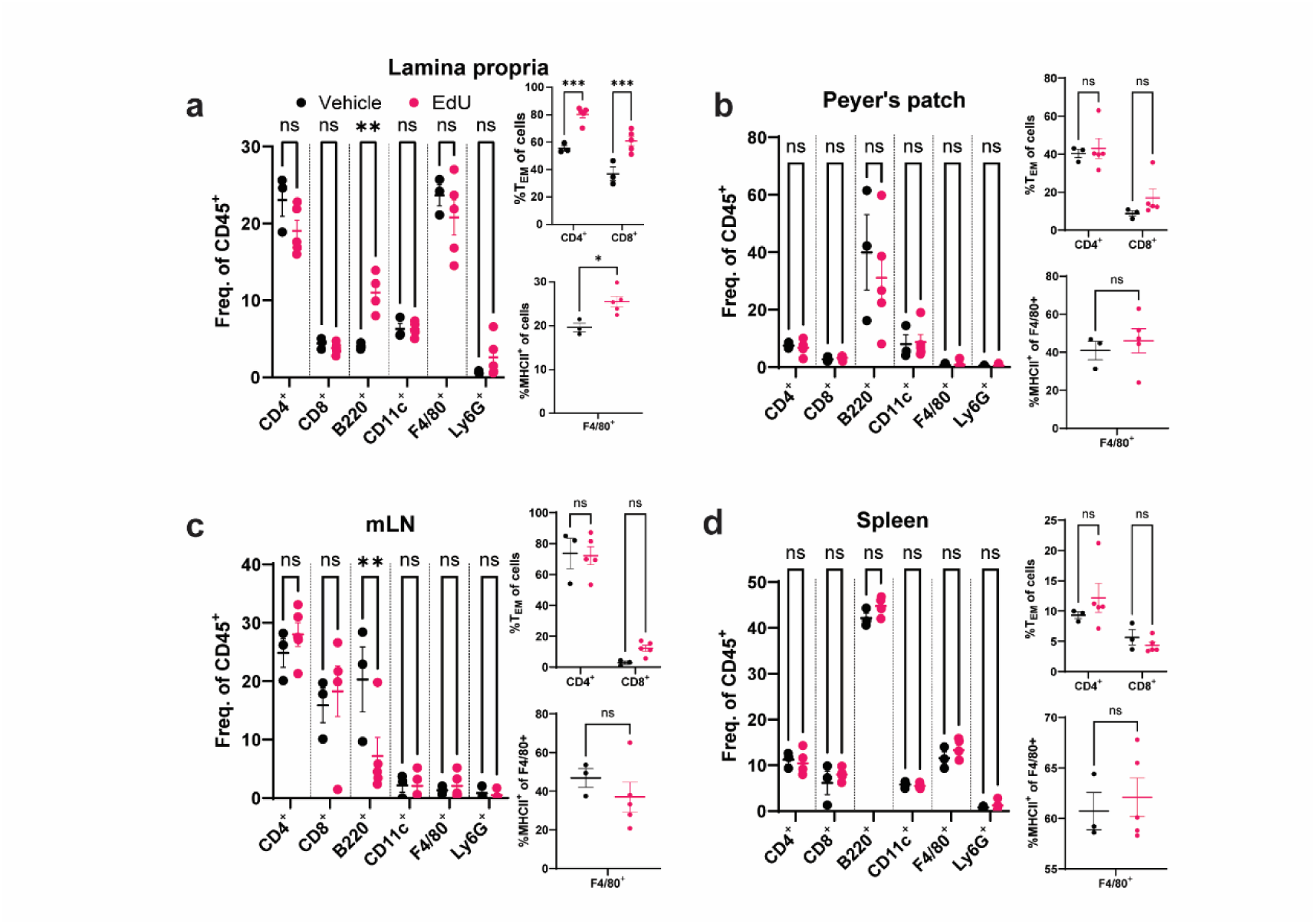
EdU exposure induces local immune system modulation. **a**,**b**,**c**,**d**, (Left) Frequencies of the indicated cell types within the CD45^+^ population of (**a**) lamina propria, (**b**) Peyer’s patch, (**c**) mesenteric lymph node, (**d**) spleen of mice that underwent 14 d of EdU. (Right) Frequency of T effector memory (T_EM_, CD62L^-^ CD44^+^) within the indicated T cell compartment and frequency of MHCII^+^ events of F4/80^+^ cells. *n* = 3-5 mice per genotype. One-way analysis of variance (ANOVA) with Tukey’s multiple comparisons post hoc test and two-tailed Student’s t test in (**a**,**b**,**c**,**d**). *p < 0.05, **p < 0.01, ***p < 0.001. ns, not significant.

**Extended Data Fig. 3 (corresponds to Fig. 2):**
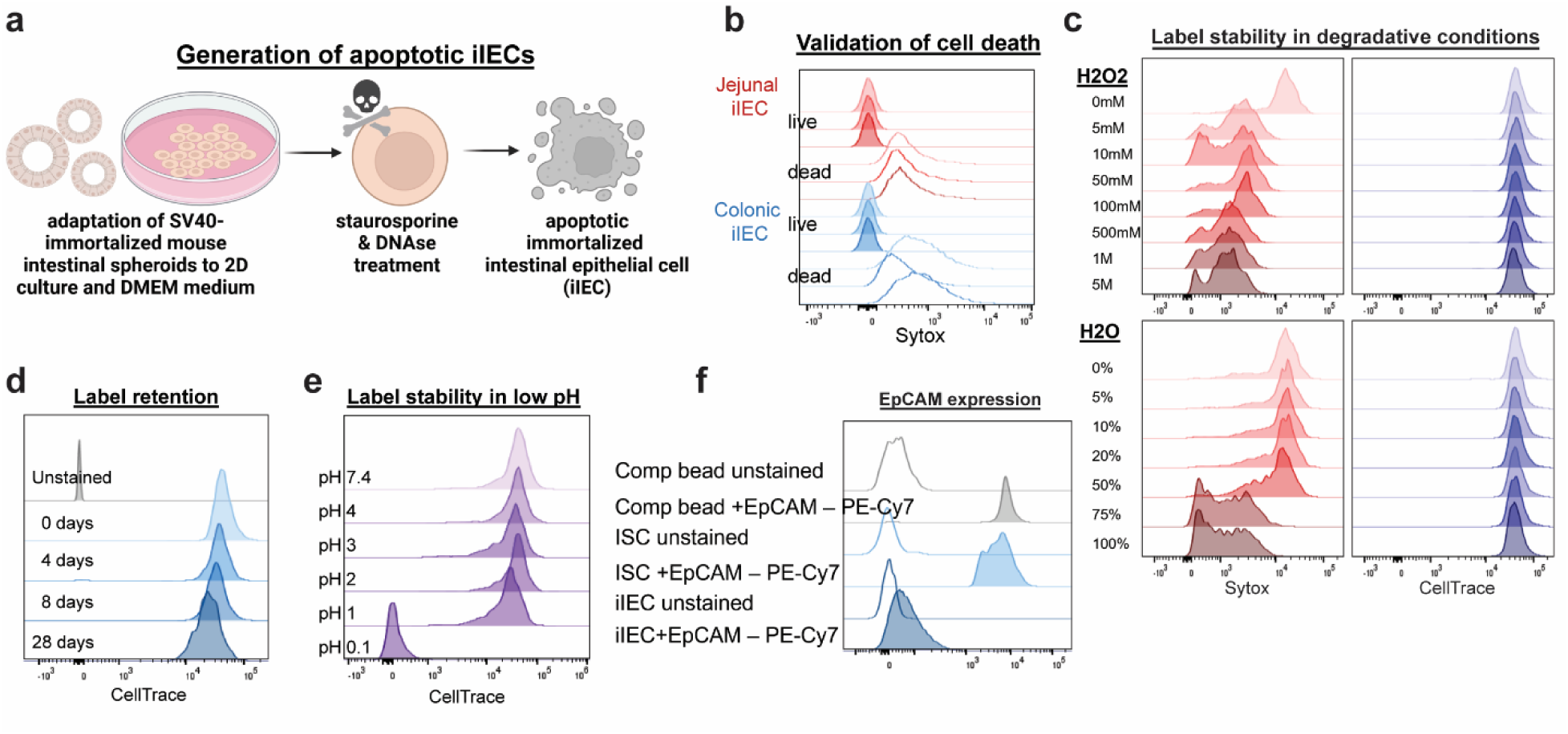
Generation of dead immortalized mouse intestinal epithelial cells. **a**, Schematic of the strategy used to generate apoptotic immortalized intestinal epithelial cells (apop.iIEC). **b**, Flow cytometric validation of apop.iIEC cell death via Sytox permeability. *n* = 3. **c**, Measurement of the stability of Sytox or CellTrace dyes in labeling apop.iIECs followed by incubation in the indicated solutions. *n* = 3. **d**, Measurement of the stability of CellTrace dye in labeling apop.iIEC at the indicated time points post storage at 4°C. *n* = 3. **e**, Measurement of the stability of CellTrace dye in albeling apop.iIECs following incubation in solutions of the indicated pH. *n* = 3. **f**, Measurement of EpCAM expression of iIECs compared to ISCs. Data are representative of at least 2 independent experiments.

**Extended Data Fig. 4 (corresponds to Fig. 2):**
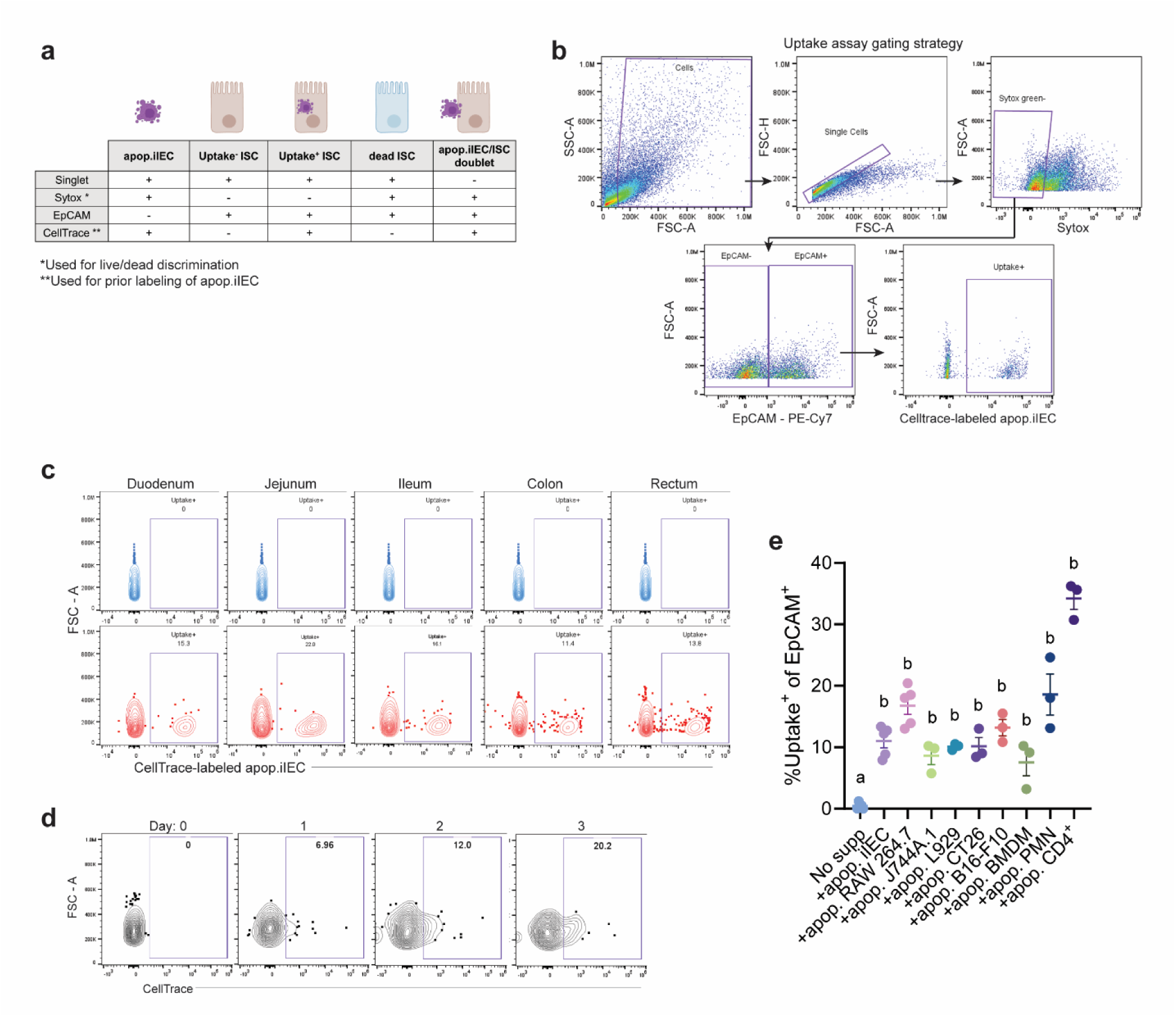
ISCs in spheroid culture uptake apoptotic cells. **a**, Schematic of the strategy used to detect apop.iIEC, uptake^+^ ISCs, uptake^-^ ISCs, dead iscs, and apop.iIEC/ISC doublets using flow cytometry. **b**, Gating strategy used to detect uptake^+^ ISCs and uptake^-^ ISCs. **c**, Representative plots of flow cytometric analysis of the frequency of uptake^+^ ISCs (Sytox^-^EpCAM^+^CellTrace^+^) spheroids grown from crypts harvested along the intestinal tract and incubated with CellTrace-labeled apop.iIECs. *n* = 3 replicates. **d**, Representative plots of flow cytometric analysis of the frequency of uptake^+^ ISCs after the indicated time points. *n* = 5 per time point. **e,** Quantification of uptake^+^ followed incubation with apoptotic cells from the indicated sources. *n* = 3 replicates. Data are representative of at least 2 independent experiments. One-way analysis of variance (ANOVA) with Tukey’s multiple comparisons post hoc test (**e**). Letters denote significant differences among groups in (**e**).

**Extended Data Fig. 5 (corresponds to Fig. 3):**
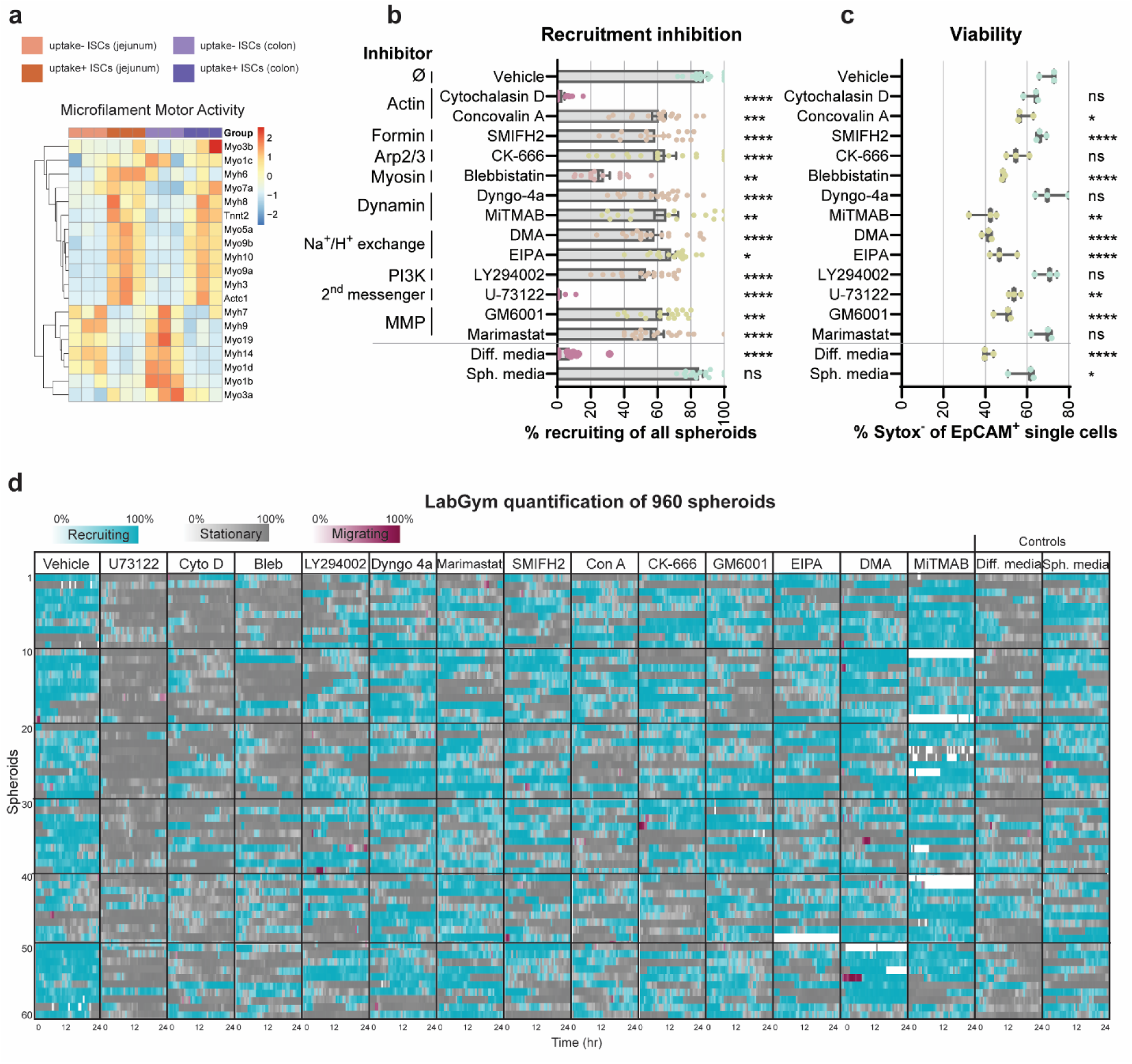
Actin-mediated spheroid behaviors decrease ISC and apoptotic cell proximity. **a**, Bulk RNAseq comparing uptake^+^ to uptake^-^ spheroids derived from mouse jejunum or colon. Microfilament motor activity gene list generated from Mouse Gene Ontology database. *n* = 3 per condition. **b**,**c**,(**b**) inhibition of recruitment and (**c**) decreased viability caused by compounds that inhibit the indicated processes. Differentiation medium and spheroid medium were used as controls. Ratio of inhibition to viability is presented. *n* = 15-18 spheroids per condition. **d**, Behavior annotation timeline of Recruiting, Stationary, and Migrating behaviors of spheroids cultured in spheroid (sph) medium or differentiation (diff) medium with the color saturation indicating the certainty of that behavior. Data are representative of at least 2 independent experiments. One-way analysis of variance (ANOVA) with Tukey’s multiple comparisons post hoc test in (**b**,**c**). *p < 0.05, **p < 0.01, ***p < 0.001, ****p < 0.0001. ns, not significant.

**Extended Data Fig. 6 (corresponds to Fig. 4):**
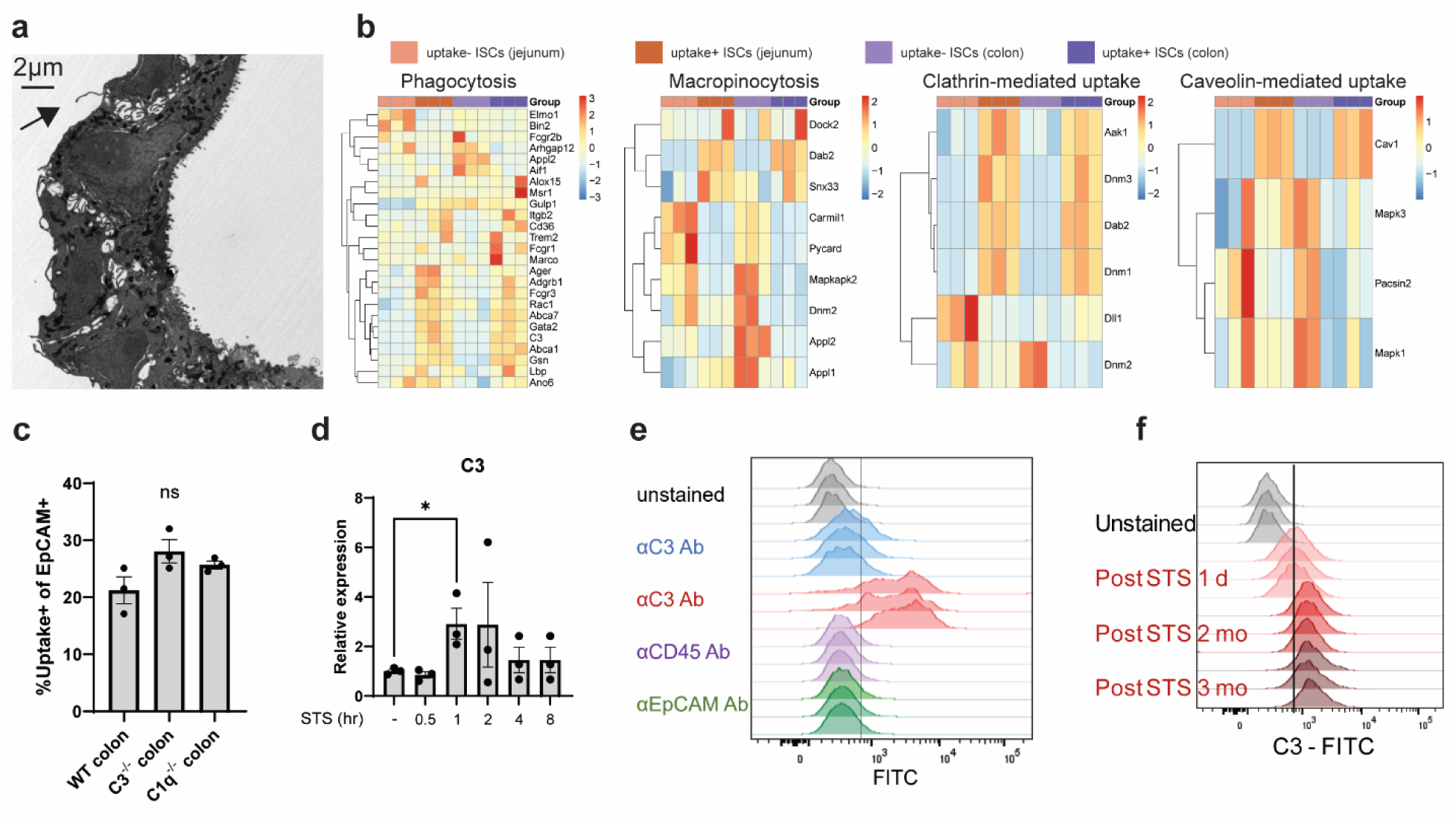
Basolateral protrusions and C3 self- opsonization of apoptotic cells underlie ISC uptake. **a**, Ultrastructure of an intact spheroid. Arrowhead denotes a basolateral membrane protrusion extended by an ISC. Scale bar = 2 µm. **b**, Bulk RNAseq comparing uptake^+^ to uptake^-^ spheroids derived from mouse jejunum or colon. Phagocytosis, micropinocytosis, clathrin-mediated uptake, and caveolin-mediated uptake gene lists generated from Mouse Gene Ontology database. *n* = 3 per condition. **c**, Frequency of apop.iIEC uptake^+^ ISCs derived from WT, C3^-/-^, or C1q^-/-^ mice. *n* = 3 per condition. **d**, Relative expression of C3 transcript after addition of staurosporine (STS) to iIEC culture with cell harvest timepoints as indicated. *n* = 3 per condition. **e**, iIEC cell surface staining using the indicated antibodies conjugated to FITC. *n* = 3 per condition. **f**, iIEC cell surface staining following addition of STS using αCD3. *n* = 3 per condition. Data are representative of at least 2 independent experiments. One-way analysis of variance (ANOVA) with Tukey’s multiple comparisons post hoc test in (**c**,**d**). *p < 0.05. ns, not significant.

**Extended Data Fig. 7 (corresponds to Fig. 5):**
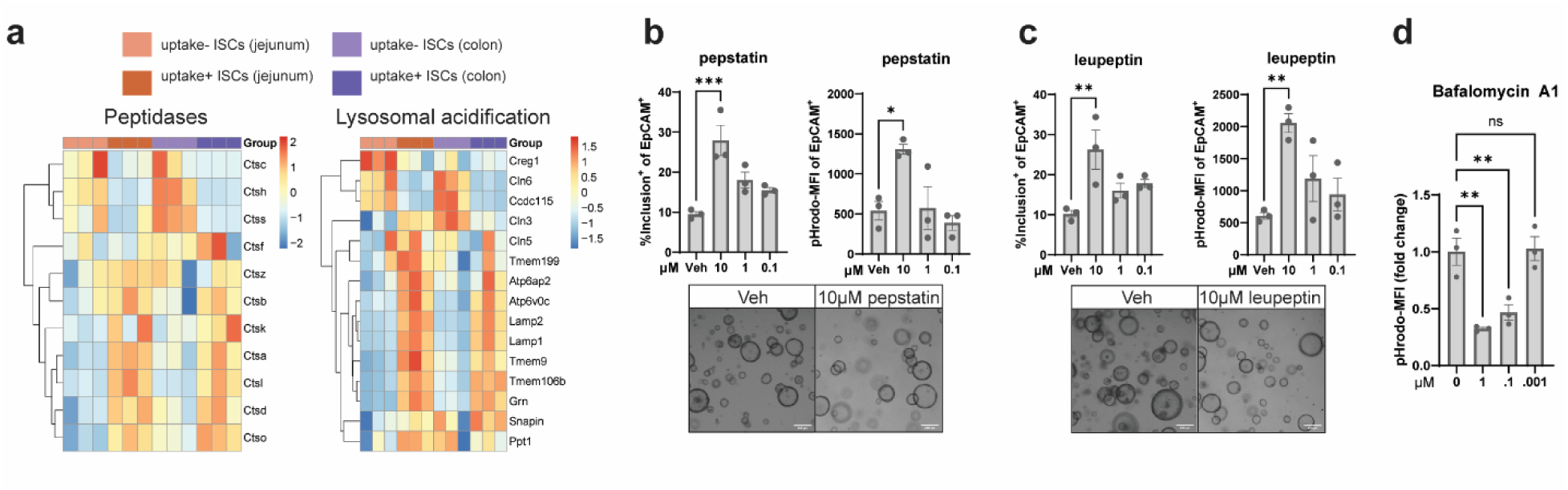
ISCs are sufficient in degrading internalized apoptotic cells. **a**, Bulk RNAseq comparing uptake^+^ to uptake^-^ spheroids derived from mouse jejunum or colon. Peptidases and lysosomal acidification gene lists were generated from Mouse Gene Ontology database. *n* = 3 per condition. **b**,**c** Percent of ISCs with an apop.iIEC inclusion and degree of acidification (pHrodo) following incubation with (**b**) pepstatin or (**c**) leupeptin. *n* = 3 per condition. **d**, Degree of acidification (pHrodo MFI) following incubation with Baflomycin A1. n = 3 for condition. Data are representative of at least 2 independent experiments. One-way analysis of variance (ANOVA) with Tukey’s multiple comparisons post hoc test (**b**,**c,d**). *p < 0.05, **p < 0.01, ***p < 0.001. ns, not significant.

**Extended Data Fig. 8 (corresponds to Fig. 6):**
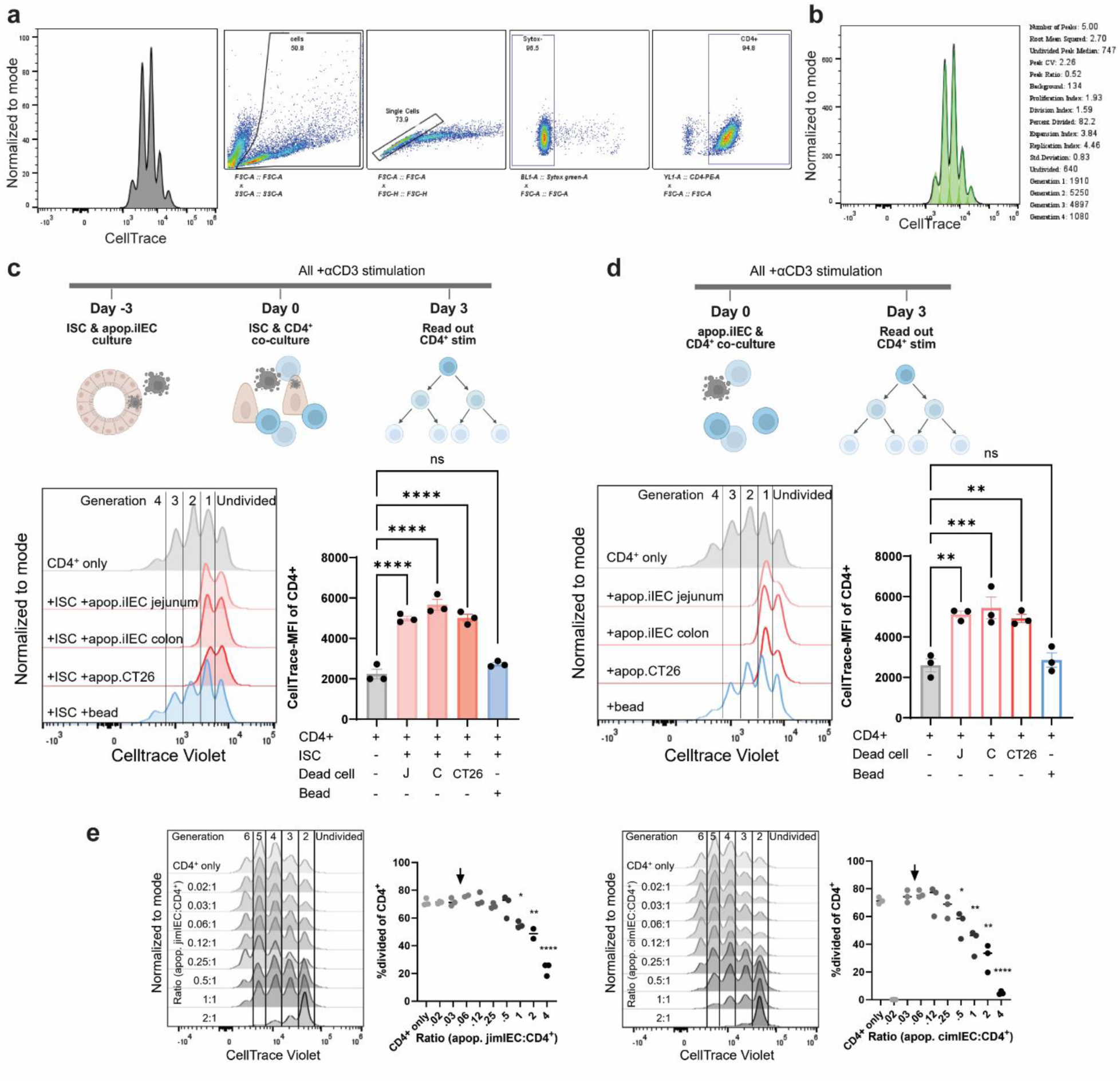
Dead cells do not contribute to enhanced CD4^+^ T cell activity in ISC and CD4^+^ co-culture assay. **a**,**b**, Gating strategy for assessing CD4^+^ T cell proliferation following co-culture with uptake^+^ or uptake^-^ ISCs in (**a**) with proliferation modeling in (**b**). **c**, Freshly isolated and CellTrace-stained splenic CD4^+^ T cells were supplemented with cells from spheroid co-culture with apop.iIECs or apoptotic CT26 cells as indicated. (Top): Schematic of experimental pipeline. (Bottom): CellTrace signal indicating CD4^+^ proliferation with quantification of CellTrace MFI. *n* = 3 per condition. **d**, Freshly isolated and CellTrace-stained splenic CD4^+^ T cells were supplemented with apop.iIECs or apoptotic CT26 cells as indicated. (Top): Schematic of experimental pipeline. (Bottom): CellTrace signal indicating CD4^+^ proliferation with quantification of CellTrace MFI. *n* = 3 per condition. **e**, Freshly isolated and CellTrace-stained splenic CD4^+^ T cells were supplemented with apop.iIECs derived from jejunum or colon in the ratios as indicated with the number of CD4^+^ T cells held constant. Arrowhead indicates the number of live uptake^+^ ISCs needed to see enhanced CD4+ T cell proliferation. *n* = 3 per condition. Data are representative of at least 2 independent experiments. One-way analysis of variance (ANOVA) with Tukey’s multiple comparisons post hoc test in (**c**,**d,e**). *p < 0.05, **p < 0.01, ***p < 0.001, ***p < 0.0001. ns, not significant.

**Extended Data Fig. 9 (corresponds to Fig. 6):**
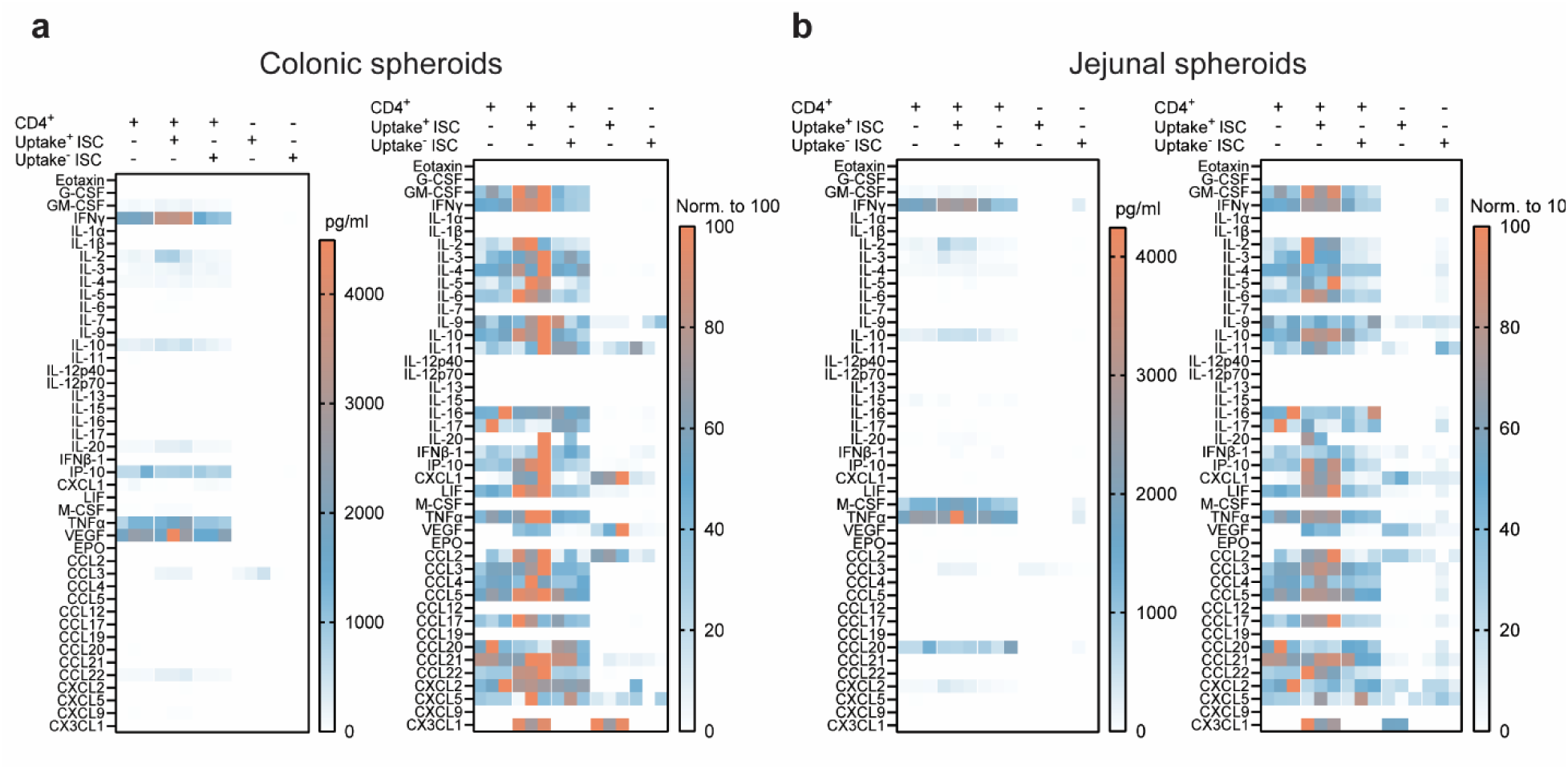
ISC uptake of apoptotic bodies enhances CD4^+^ T cell production of cytokines. **a**,**b**, Luminex cytokine analysis of ISC and splenic CD4^+^ T cell co-culture supernatant following 72 h of co-culture. (**a**) ISCs were derived from mouse jejunum. (**b**) ISCs were derived from mouse colon. *n* = 3 for all conditions.

**Extended Data Fig. 10 (corresponds to Fig. 6):**
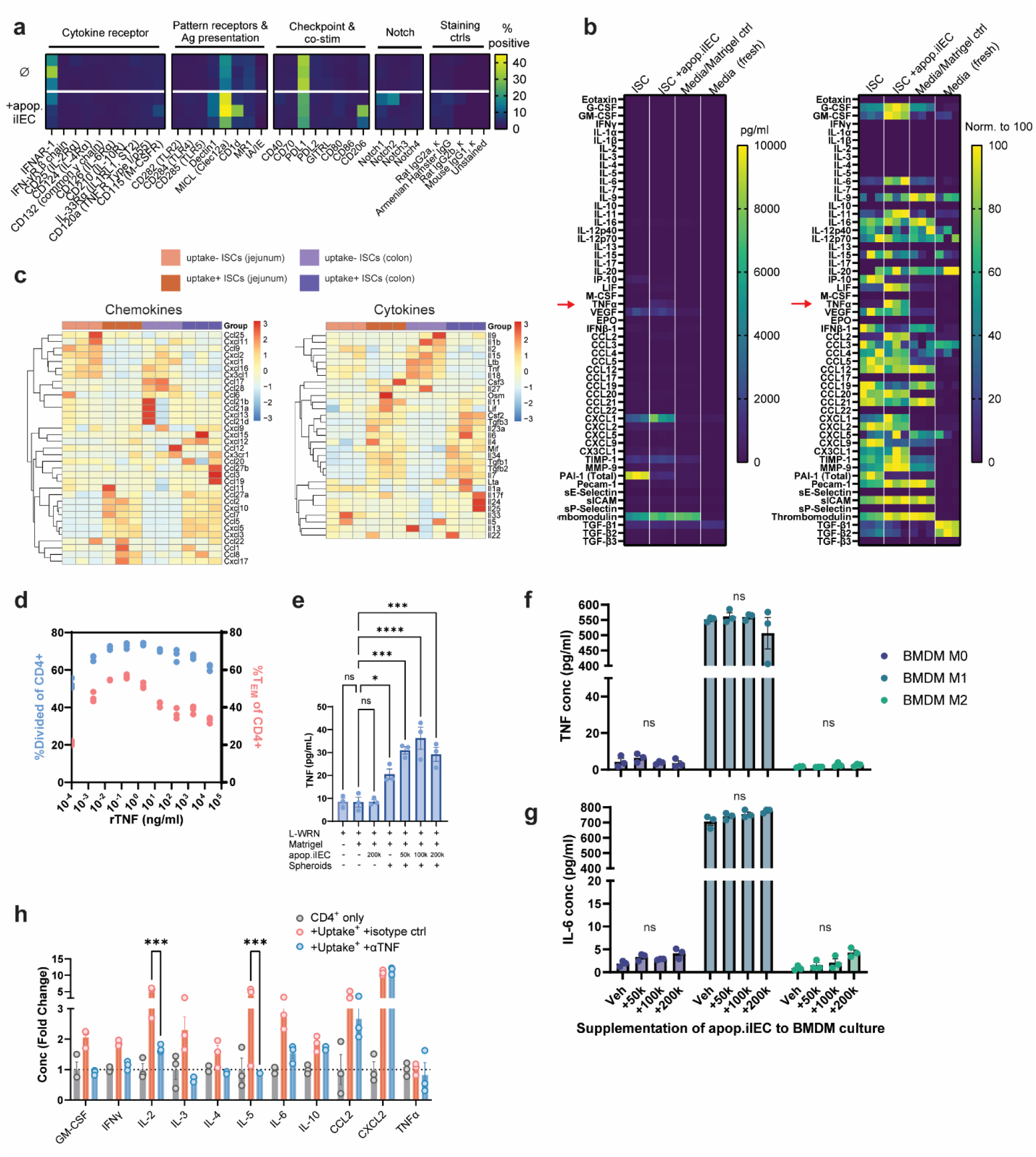
ISC apoptotic body uptake elicits T cell- enhancing activity. **a**,**b** Spheroids were cultured with or without apop.iIEC for 72 h and (**a**) underwent cell surface molecule expression by percent positive of all ISCs and (**b**) Luminex cytokine analysis of culture supernatant. *n* = 3 for all conditions. **c**, Bulk RNAseq comparing uptake^+^ to uptake^-^ spheroids derived from mouse jejunum or colon. Chemokines, cytokines, and TNF target genes were from KEGG database. *n* = 3 per condition. **d**, Freshly isolated and CellTrace-stained splenic CD4^+^ T cells were supplemented with recombinant TNF (rTNF). Cell division (CellTrace signal) and %T_EM_ (CD62L^-^CD44^+^) were assessed by flow cytometry. *n* = 3 for all conditions. **e**, ISCs were cultured (200k per well) with the indicated numbers of apop.iIECs in 50% L-WRN for 72 h. TNF concentration of the culture supernatant was measured by ELISA. *n* = 3 for all conditions. **f**,**g**, Bone marrow-derived macrophages (200k per well) were supplemented with polarizing cytokines (LPS and IFNγ for M1, IL-4 for M2) and the indicated numbers of apop.iIECs for 24 h. (**f**) TNF and (**g**) IL-6 concentration of the culture supernatant was measured by ELISA. *n* = 3 for all conditions. Data are representative of at least 2 independent experiments. One-way analysis of variance (ANOVA) with Tukey’s multiple comparisons post hoc test (**e**,**f**,**g**,**h**). *p < 0.05, ***p < 0.001, ***p < 0.0001. ns, not significant.

